# The non-catalytic ε DNA polymerase subunit is an NPF motif recognition protein

**DOI:** 10.1101/2025.03.17.643635

**Authors:** Salla Keskitalo, Boglarka Zambo, Antti Tuhkala, Kari Salokas, Tanja Turunen, Norbert Deutsch, Norman Davey, Zsuzsanna Dosztányi, Markku Varjosalo, Gergo Gogl

## Abstract

Short linear motifs (SLiMs) in disordered protein regions direct numerous protein–protein interactions, yet most remain uncharacterized. The Asn-Pro-Phe (NPF) motif is a well-known EH-domain ligand implicated in endocytosis, but here we reveal that the non-catalytic subunit of human DNA polymerase ε (POLE2) also serves as a general NPF-motif receptor. Using a quantitative “native holdup” assay, we find that POLE2 selectively binds diverse NPF-containing peptides, including canonical EH-domain ligands (e.g., SYNJ1) and previously uncharacterized motifs. Biochemical measurements and mutational analysis show that NPF motifs interact with a shallow pocket near the POLE2 C-terminus, and AlphaFold predictions confirm key roles for Y513, E520, and S522 in motif coordination. Proteome-scale affinity screens identify NPF-containing nuclear proteins (e.g., WDHD1, DONSON, TTF2) that bind POLE2 with micromolar affinities, and their motif mutations abolish binding in cell extracts. Although POLE2 primarily tethers the catalytic POLE subunit to replication forks, these results indicate that it can also recruit various NPF-bearing partners involved in replication, DNA repair, and transcription regulation. Notably, NPF motifs optimized for EH-domain binding can still associate with POLE2, highlighting the inherent degeneracy of SLiM-mediated networks. Overall, these findings establish POLE2 as a central hub linking replication with other processes via broad NPF-motif recognition.

## Introduction

Regions showing strong local evolutionary conservation frequently occur in otherwise poorly conserved, intrinsically disordered protein regions. Many of these mediate protein-protein interactions and these sites are often referred to as SLiMs (for **s**hort **l**inear **i**nteraction **m**otifs) or simply as motifs ^1,2^. Motifs are indispensable for complex biological processes and constitute the majority of our cellular interactome generating a great interest in identifying novel motifs and motif recognition proteins. To this end, several computational *de novo* motif discovery tools have been developed to identify putative motifs in disordered protein regions based on their unique amino acid compositions and evolutionary properties ^3,4^. Although these approaches identify numerous putative motifs, they cannot predict their binding partners, leaving them as orphan motifs. Finding their unknown partners poses significant experimental challenges, since motif-mediated interactions typically have weak affinities and fast binding kinetics ^5^. These properties make them practically invisible for mainstream interactomic approaches. Fortunately, it is possible to exploit the degenerate nature of motif-mediated interactions to predict the putative binding partner of a given orphan motif in certain cases. By comparing the sequence of an orphan motif to catalogs of consensus sequences of known motif-recognizing proteins, it is sometimes possible to assign the putative motif to a given interaction partner ^6^. However, a simple consensus motif can be often recognized by multiple members of an entire protein family ^7^, or members of multiple protein families ^8^ and simplistic consensus motifs do not contain sufficient information to distinguish between interactions of different strength ^9^. Adding to these limitations, we are currently unaware of how many motif recognizing proteins may be present in our proteome and only a few hundreds of consensus motifs have been cataloged so far ^10^.

The case of NPF (Asn-Pro-Phe) motifs illustrates these critical issues. Almost three decades ago, it was discovered that NPF motifs can bind to EPS15 homology (EH) domains ^11–13^ (Figure S1A). EH domains are calcium-binding EF-hand domains found in eleven human EH proteins ^14^ (Figure S1B). Two of these EH proteins contain three EH domains, four of them contains two domains and the four EHD proteins that contains a single EH domain form stable oligomers ^14,15^. The partners of these proteins contain disordered protein regions containing NPF motifs, often present in multiple copies, contributing to multivalent and synergistic interactions ^16^. EH proteins primarily function during vesicle formation, and their NPF motif-mediated interactions are crucial for protein sorting ^16,17^. These interactions exploit the stable Asx turn conformation formed by the NPF tripeptide, where the carbonyl of the Asn sidechain forms a hydrogen bond with the amide nitrogen of Phe residue of the NPF motif ^13^. Since all members of the EH family recognize the same structural motif, a putative NPF motif can interact with any members of the family. In addition, many additional proteins have been reported to be able to bind NPF motifs, that are unrelated to EH domains (Figure 1A). NPF motifs are also involved in vesicle budding interactions of SNAREs through mediating interactions with the SEC23/24 subunits of COPII ^18,19^. The NPF motif of Pygo2 can be recruited at the composite interface formed between an SSBP2 dimer and LDB1 and their ternary complex forms the so called WNT enhanceosome ^20,21^. An N-terminal NPF motif in BORA was found to be part of a critical region for activating Aurora A kinase (AURKA), and it was proposed that this motif may interact with the AURKA kinase domain ^22^. A fungal protein called SHD1 binds to NPF motifs that functions as endocytic internalization motifs ^23^. In a phage display experiment, the HIV-1 envelope glycoprotein gp41 was found to have a strong preference for binding HxxNPF motifs and such peptides were able to inhibit HIV-1 envelope glycoprotein-mediated syncytium-formation ^24^. Finally, four studies were published in parallel to our investigation, two reporting that the non-catalytic ε DNA polymerase subunit (POLE2) can interact with NPF motifs found in Protein downstream neighbor of Son (DONSON) ^25,26^, and two reporting an interaction between POLE2 and the transcription termination factor 2 (TTF2) ^27,28^. Besides these NPF binding proteins, our proteome contains many additional Pro-rich motif recognizing proteins – like SH3 domains that bind to PxxP motifs or PTB domains that bind to NPxY motifs – and their binding motifs may also partially overlap with NPF motifs ^29^. As illustrated above, a putative NPF-containing motif can potentially bind to large numbers of partners, including the 11 type EH proteins, COPII, SSBP2/LDB1, AURKA, POLE2, and possibly many others including proteins from pathogens. Currently, it is not yet explored what are the determinants of specificity between these different types of interactions and it is also unknown how widespread are the interactomes of each of these NPF recognition proteins.

**Figure 1.**
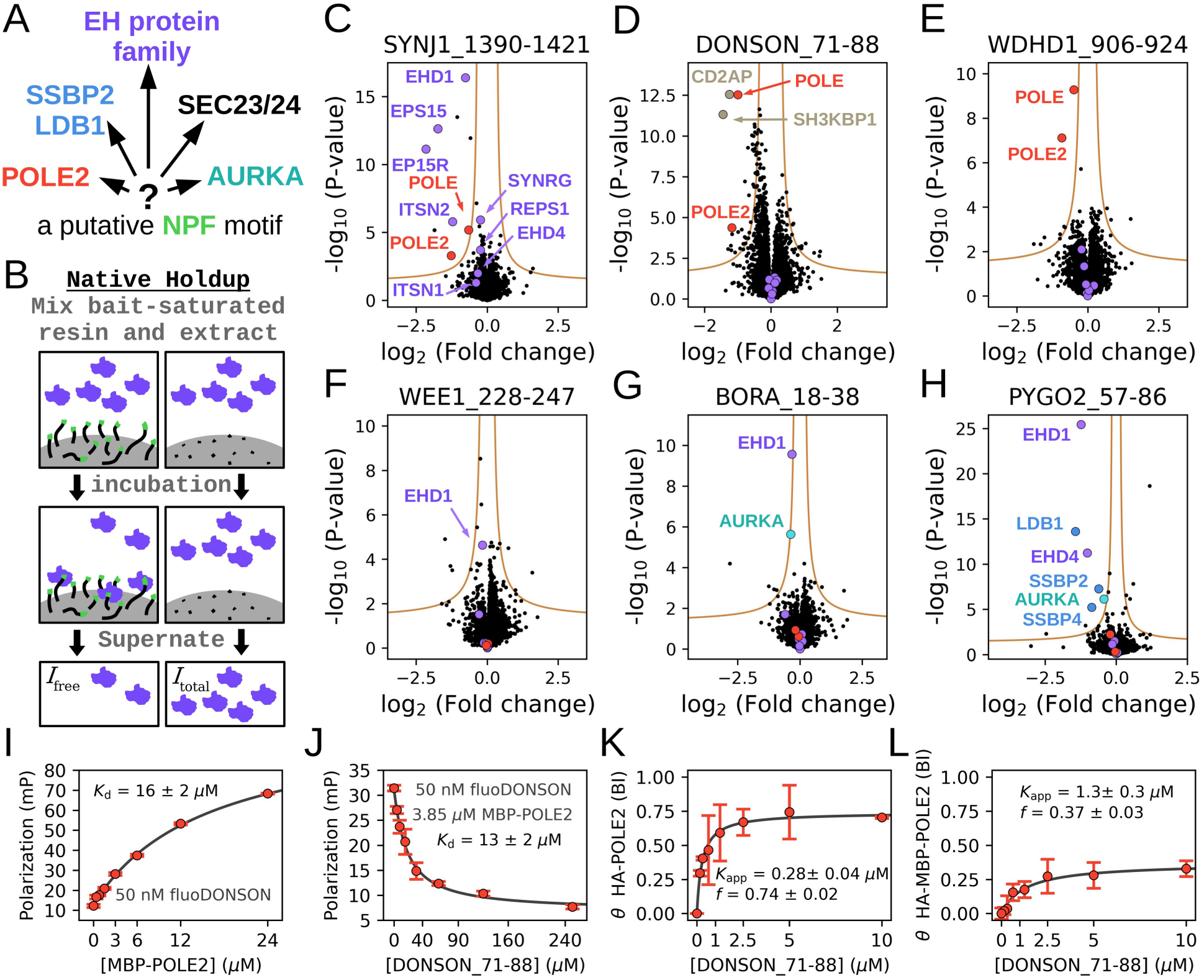
NPF motifs bind to various types of proteins, including POLE and POLE2. (A) NPF tripeptides were described to interact with multiple types of receptors, making prediction of partners of putative motifs highly ambiguous. (B) The native holdup (nHU) affinity interacotmic approach is ideal to capture motif-mediated interactions because of high ligand concentration allowing abundant complex formation of weak interactions and no washing procedures that would eliminate transient interactions. (C-H) Six different NPF motif peptides were assayed with nHU coupled with mass spectrometry. Interaction partners belonging to known or suspected NPF motif receptors are highlighted. Experiments were carried out with 2 biological and 3 technical replicates (total N = 6). (I) Fluorescence polarization was measured by titrating recombinant avi-His_6_-MBP-POLE2 to a fluorescent NPF motif peptide, confirming binary complex formation. (J) Competitive titration experiments show that the non-fluorescent NPF motif peptide can compete with the binding of the labeled one with similar affinity. *In vitro* binding experiments were carried out in triplicates. (K) *Ex vivo* measured affinity of HA-POLE2, measured by nHU in cell extract, shows higher affinity than *in vitro* measured one, measured with purified recombinant protein. (L) The affinity of HA-MBP-POLE2 was found to be weaker than of HA-POLE2 with a clear partial binding activity. Titration nHU experiments were carried out with at least two technical replicates. See Table S1, S2, Figure S2, S3, S4, S5 for more details.

Here, we developed an experimental pipeline to identify partners of NPF motifs and found that POLE2 is a general NPF motif recognition protein. We unraveled a proteome-wide NPF motif-mediated interaction network of POLE2, we investigated their binding mechanism and determined the key specificity determinants of NPF motifs between POLE2 and an EH protein, called EPS15. Our findings provide new insights into the broad interaction network of DNA polymerase ε and its role in the replisome. Furthermore, this study highlights the fundamental specificity issue in motif-mediated interactions and presents a comprehensive, unbiased experimental framework to address it.

## Results

### Identification of POLE/POLE2 as a recurring interaction partner of unrelated NPF motif-containing peptides

In order to identify the partners of NPF motifs, we used the native holdup (nHU) assay that was recently developed by our team (Figure 1B). The holdup approach has been extensively used to determine equilibrium dissociation constants of transient interactions mediated by PDZ and SH3 domains, 14-3-3, and human papilloma virus E6 proteins, and even nucleic acid-protein interactions ^30,31,7,32–35^. In the nHU experiments, a purified bait molecule, or a control compound, is immobilized on a resin at high (typically 10 μM, or higher) concentration and these resin stocks are subsequently mixed with dilute cell extracts. After a relatively long incubation, i.e. 2 h at 4 °C, the liquid phase is separated by centrifugation. This liquid phase contains the unbound prey molecules at near binding equilibrium that can be quantified using selective and quantitative analytical approaches, such as mass spectrometry. Therefore, proteins that interact with the bait molecule will appear to be depleted in the solvent phase and by measuring its relative concentration compared to the control nHU experiment, the degree of binding (*θ*) can be determined that can be converted to apparent steady-state dissociation constants, according to:

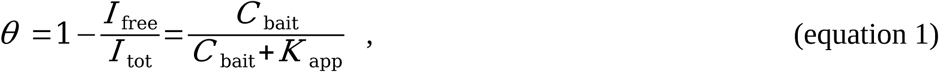

where *I*_tot_ and *I*_free_ are the measured total and bait-depleted equilibrium mass spectrometry total ion intensities of prey proteins, *C*_bait_ is the bait molecule concentration and *K*_app_ is the apparent dissociation constant.

To demonstrate that nHU could be used to characterize interactions of NPF motifs, we selected three previously reported NPF motifs from SYNJ1, BORA and PYGO2, that are expected to bind to EH proteins, AURKA and SSBP2/LDB1, respectively. We also included three orphan or poorly studied NPF motifs from DONSON (likely to bind to POLE2), WDHD1 and WEE1 that were predicted by the SLiMPrints approach over a decade ago ^4^. We synthesized these motifs as biotinylated peptides to use them in nHU experiments as baits using biotin in the control experiment. Subsequently to nHU assay, we analyzed the depleted extracts with label free quantitative mass spectrometry (MS) and identified statistically significant interactions as previously described ^35^ (Figure 1C-H, Table S1). As expected, the SYNJ1 NPF-motif rich region – containing three NPF motifs – showed significant interaction with EH proteins EHD1, EPS15, EP15R, ITSN2 and showed substantial depletion, albeit below our stringent significance threshold in the EH proteins SYNRG and REPS1. The NPF motif of WEE1 depleted EHD1, albeit the statistical significance of this interaction was below our strict threshold. The motif of BORA also depleted EHD1, as well as AURKA. PYGO2 was found to interact with both EHD1, EHD4, and AURKA, besides the core WNT enhanceosome, consisting of LDB1 and SSBP2/4. The NPF motif of DONSON showed significant interaction with two SH3 domain-containing proteins, CD2AP and SH3KBP1. Most surprisingly, the NPF motif peptides of both DONSON, WHDH1 and even SYNJ1 showed significant binding to the hetero-dimer of catalytic and the non-catalytic subunits of the ε DNA polymerase, called POLE and POLE2, respectively. Although the POLE/POLE2 complex was already suspected to interact with DONSON through its NPF motif, it was rather unexpected to identify multiple types of evolutionarily unrelated NPF motifs as binding partners of same complex, including the well characterized EH domain-binding NPF motif from SYNJ1. To confirm these findings, the proteome-wide nHU experiments coupled to MS were repeated with the DONSON, WDHD1 and SYNJ1 peptide baits with consistent results and with the additional observation of an interaction between WDHD1 and EHD4 (Figure S2A). The interaction between these distinct NPF motifs and cellular POLE was also validated with targeted Western-blot analyses (Figure S2B, S3).

### Direct interaction between NPF motifs and POLE2

The experiments revealed that POLE2 consistently showed higher depletion than POLE in binding studies, indicating that the NPF motif directly interacts with POLE2 and captures POLE indirectly through a reversible POLE-POLE2 interaction (Figure S4). *In vitro* binding experiments confirmed this direct interaction. Recombinant POLE2, tagged with His_6_-avi-MBP, was tested with a fluorescein-labeled DONSON_71-88 peptide in fluorescence polarization assays, yielding a dissociation constant of 16 μM (Figure 1I). Competitive binding assays using a non-fluorescent DONSON_71-88 peptide corroborated this result, with a dissociation constant of 13 μM (Figure 1J).

In cellular contexts, nHU titration experiments with HA-POLE2 and DONSON_71-88-saturated resin demonstrated an apparent affinity of 0.28 μM, significantly stronger—approximately 40-fold—than the *in vitro* affinity to MBP-tagged POLE2 (Figure 1K, S5, Table S2). This disparity is likely due to the destabilization of POLE2 when isolated from POLE and the influence of the MBP tag. Further experiments with HA-MBP-POLE2 in HEK293T cell extracts revealed 10-fold weaker binding compared to HA-POLE2 (Figure 1L, S5, Table S2). HA-MBP-POLE2 also exhibited partial activity where only one out of three molecules could bind the NPF motif.

In summary, despite variations in apparent binding affinity caused by destabilization or the MBP tag, the experiments demonstrated that POLE2 binds directly and reversibly to the NPF motif of DONSON, even in the absence of POLE.

### Structural insight into NPF motif binding by POLE2

We used AlphaFold 3 (AF3) to predict the structure of NPF motif-bound POLE2 ^36^ (Figure 2). Based on these predictions, the NPF motif of DONSON is bound to POLE2 by adopting a compact Asx turn conformation, similarly to NPF motifs are bound to EH domains or the SSBP2/LDB1 complex. This core NPF motif is bound to a shallow pocket of POLE2, close to its C-terminus. E520 and S522 from the last beta strand of POLE2 are coordinating the amide nitrogen of the Asn sidechain of the NPF motif. The aromatic Phe of the NPF is involved with several hydrophobic contacts, including Y311, that is also in potentially hydrogen bond distance to the carbonyl of the Asn carboxamide side-chain of the NPF motif. Beside the core NPF motif, the N-terminal flanking region also has both main- and side-chain mediated interactions. Y513 from the penultimate beta strand of POLE2 interacts with the amide nitrogen of Asn of the NPF motif. Finally, D284 forms a salt bridge with the NPF motif preceding Arg residue of DONSON.

**Figure 2.**
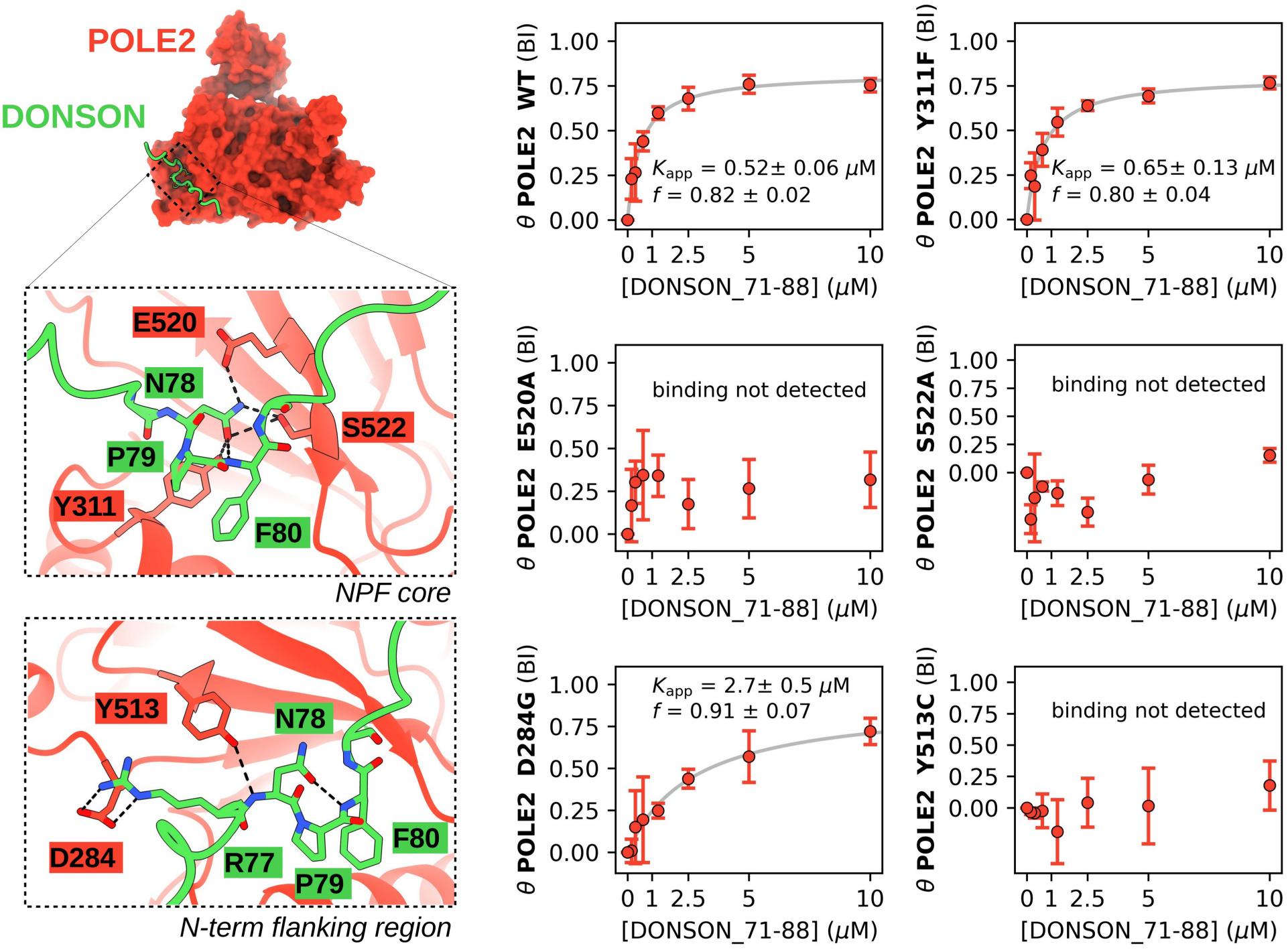
NPF motifs bind to a specific interface on POLE2. (left) AlphaFold predictions confidently dock NPF peptides onto a solvent-exposed surface of POLE2 found near its C-terminus. Two insets show directed interactions of the core NPF tripeptide and its N-terminal flanking region. (right) In order to interfere with NPF motif binding, POLE2 residues Y311F, E520, and S522 were selected to be mutated to Ala. In addition, the rare, but natural D284G and Y513C mutations were introduced that is predicted to interact with the N-terminal flanking region of NPF motifs. Titration nHU experiments carried out with these HA-POLE2 point mutants confirms the predicted bound conformation. Titration nHU experiments were carried out with at least two technical replicates. See Table S2, Figure S6, S7 for more details.

To validate these predictions, we introduced Y311F, E520A, S522A artificial mutations in HA-POLE2 to disrupt specific interactions of the core NPF motif and expressed these variants in 293T cells as well. We also introduced D284G, Y513C mutations that are uncharacterized rare, but naturally occurring polymorphic variants of POLE2 found on the putative NPF motif-binding interface. Then, prepared cell extracts from each variant and confirmed that none of these mutations caused neither decreased expression of POLE, POLE2, nor degraded products (Figure S6A-B). We performed titration nHU experiments using the DONSON NPF motif as a bait and used dot blot with an antibody against the HA tag to quantify POLE2 depletion (Figure 2, S7, Table S2). This experiment revealed that E520A, S522A, as well as Y513C completely disrupted the interaction and no residual binding could be observed with these variants, confirming that the NPF motif is coordinated by multiple residues from the last and the penultimate beta strands of POLE2. In contrast, the D284G mutation only decreased NPF motif binding by 5-fold and the Y311F mutation did not have any significant impact, showing that Y311 is not contributing to the NPF motif binding via its hydroxy group and that the N-terminal flanking region of the NPF motif can potentially interact with D284 but is not essential for binding. Importantly, both natural POLE2 variants decreased NPF motif binding, classifying D284G as a partial loss of function and Y513C as a complete loss of function variant for NPF motif binding. Further investigations are needed to evaluate the clinical importance of these natural variants, or similar POLE2 mutations.

### POLE2 interacts with NPF motifs proteome-wide

So far, our study only focused on a few hand-picked NPF motifs and studied them as short synthetic peptides. However, many other proteins may be present in the proteome that could bind to POLE2 and their putative NPF motifs are embedded in their natural structures that can greatly affect their binding properties. To investigate interactions between POLE2 and intact, full-length proteins proteome-wide, we used nHU-MS experiments with recombinant His_6_-avi-MBP-tagged POLE2 as bait and purified His_6_-avi-MBP as a control. Even though we are aware that this recombinant POLE2 may bind to NPF motifs with a somewhat altered affinity, we assumed that this will affect all NPF motif-mediated interactions equally, allowing an unbiased affinity ranking of all identified partners. Single point nHU measurements were performed at 26 μM POLE2 concentration and the recovered supernate was measured by label free proteomic MS measurements (Figure 3A, S8A, S9, Table S1).

**Figure 3.**
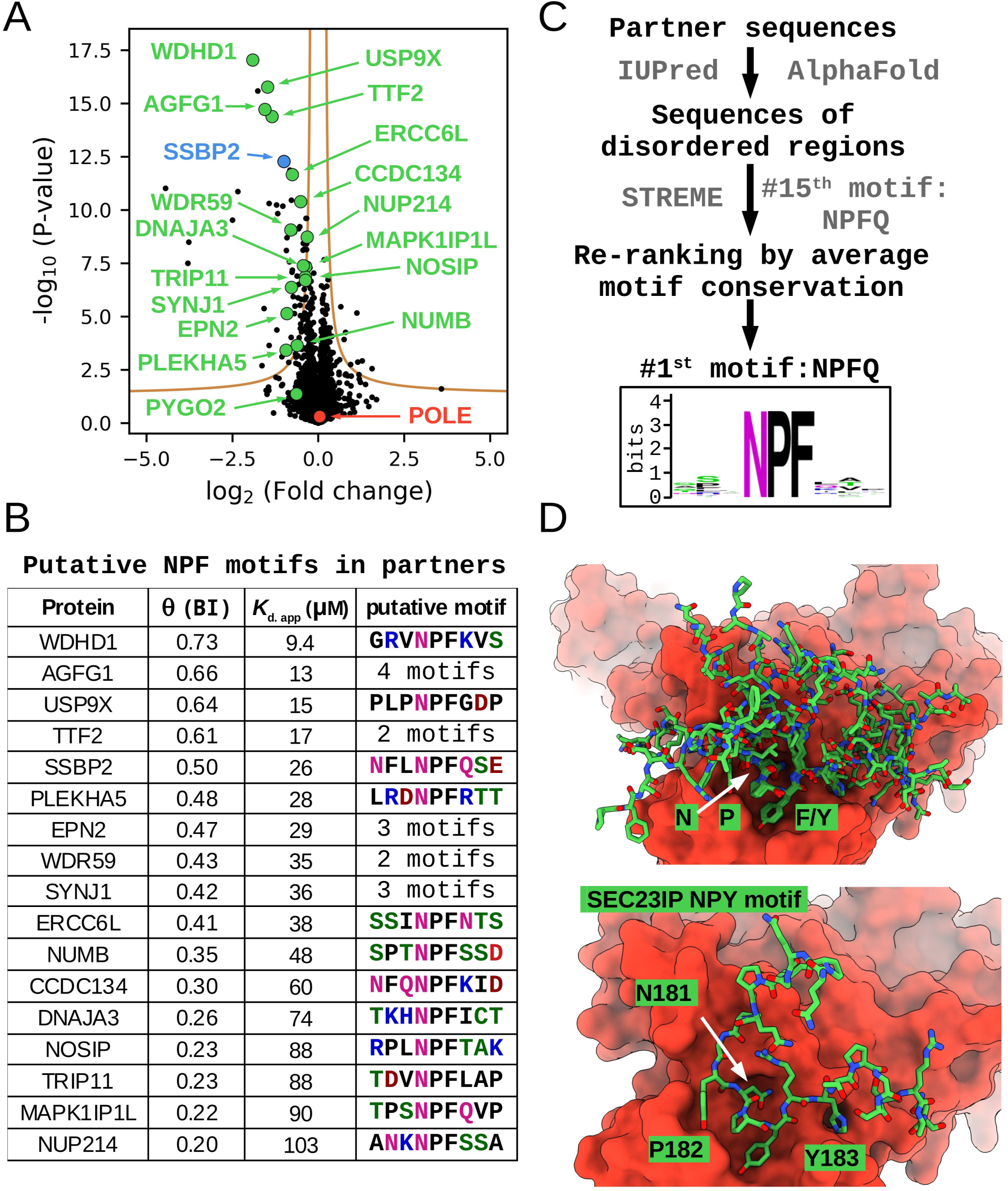
POLE2 binds to NPF motifs proteome-wide. (A) Recombinant avi-His_6_-MBP-POLE2 was used as a bait in a nHU experiment coupled with mass spectrometry revealing numerous binding partners. Experiments were carried out with 2 biological and 3 technical replicates (total N = 6). Those significantly depleted partners that contain putative NPF motifs are highlighted with green, or blue circles. (B) List of partners with putative NPF motifs. (C) Schematic pipeline of our *de novo* motif discovery approach. Based on the sequences of the identified partners, NPF motifs are the most evolutionary conserved ones out of all enriched motifs. (D) AlphaFold3 predictions reveal that although most partners bind via NPF motifs, the same binding interface is also compatible with a putative NPY motif of SEC23IP. See Table S1, Figure S8, S9, S10, S11 for more details.

In this experiment, we performed an affinity measurement with 6516 endogenous, full-length proteins, covering 32 % of the human proteome. Out of these detected proteins, 70 showed significant binding to POLE2. Interestingly, no POLE binding was detected during the experiment, possibly due to the competition with endogenous, untagged POLE2 or due to the presence of the N-terminal tag. Nevertheless, the detected full-length partners of POLE2 included SYNJ1, NUMB, EPN2 and AGFG1, well known binding partners of EH domains. In addition to these proteins with known NPF motifs, the screen identified WDHD1 as the most significant POLE2 interaction partner, whose isolated NPF motif was studied above. The binding of TTF2 was also identified with high significance, which is a recently reported NPF motif-mediated partner of POLE2. To find other partners containing NPF motifs, we used the SLiMSearch algorithm to find putative NPF motifs in disordered regions of the POLE2 partners and identified 17 proteins with such motifs (Figure 3B). The NPF consensus sequence is very short, and due to the lack of additional constraints, it is relatively common in the proteome, accounting for 509 NPF motifs in disordered regions of 446 full-length proteins out of the approximately 20k human proteins ^37^. Based on this, 70 randomly selected proteins are expected to contain only 1 or 2 NPF proteins. In contrast, in our experiments 17 POLE2 partners contained NPF motifs, showing a >10-fold enriched in proteins containing NPF motifs in the observed list of interaction partners.

It was previously found that EH domains can bind not only NPF motifs but also to related DPF, GPF, and NPY motifs ^38,39^. To investigate the possibility of similar interactions with POLE2, we explored if de novo motif discovery tools, such as STREME ^40^ or SLiMFinder ^41^ were able to identify recurring motif sequences in the identified POLE2 partners. Unfortunately, these tools failed at this task. STREME mostly identified structural motifs or repeats in the sequences of the identified POLE2 partners, while SliMFinder ^41^ did not discover any meaningful motif. Using the sequences of only the disordered regions of these partners the STREME algorithm could identify the NPFQ consensus motif but only as the 15th most enriched motif type (Figure S10A, Table S3). Then, we calculated an evolutionary conservation score for each motif instance identified in the STREME output and re-ranked the proposed motifs based on the average evolutionary score of the motifs belonging to the same type (see Material and Methods) (Figure 3C, S10B, Table S3). This highlighted the NPF consensus motif which was the most conserved out of all.

This approach identified 19 motifs in 14 proteins that belonged to this NPFQ motif type. All of them contained the core NPF tripeptide sequence, but they had diverse residues at the last position. No related motif classes, such as DPF or NPY, were identified by this approach. We also carried out binary AF3 predictions, using the full-length sequences of POLE2 and its 70 significant interaction partners. This approach identified seven partners that bind to this particular surface of POLE2, including all four proteins that were well-studied partners of EH proteins, as well as WDHD1 and TTF2 (Figure 3D, S11). Interestingly, AF3 also predicted interaction with the SEC23-interacting protein (SEC23IP) to the same surface of POLE2 but with an NPY motif. Although this protein is localized to the endoplasmic reticulum and thus spatially separated from POLE2, their intrinsic interaction indicates that the POLE2 binding pocket is compatible with NPY motifs and future studies may also identify similar motifs in other partners.

### Validation of selected nuclear POLE2-NPF motif interactions

Beside the critical role of POLE/POLE2 in replication, POLE is also known to be involved in proofreading, nucleotide excision repair and to a lesser extent in mismatch repair ^42^. Similarly, among the identified interaction partners of POLE2, only WDHD1 and TTF2 are known to be involved in replication control, yet many other partners are involved in other nuclear processes, such as transcription regulation or DNA repair. Out of these partners, we selected for further studies WDHD1, TTF2, and ERCC6L. WDHD1 is one of the most significant partners of POLE2 and its NPF motif interacts with POLE/POLE2 with high selectivity. TTF2 – a protein that terminates transcribing RNA polymerase II from the DNA template and that seems to participate in replisome disassembly – contains 2 putative NPF motifs, of which AF3 docked one to POLE2 in binary structure predictions. The DNA excision repair protein ERCC-6-like (ERCC6L) DNA helicase contains a single putative NPF motif. The cellular role of ERCC6L is rather understudied and the detected interaction between POLE2 and ERCC6L may be also important for polymerase recruitment during excision repair.

To further investigate these partners, we first synthesized their putative NPF motifs and performed nHU titration experiments against HA-POLE2 taken from cell extracts (Figure 4A, S12, Table S2). This experiment revealed that the NPF motif of WDHD1 displays a relatively weak interaction with POLE2. Out of the two putative NPF motifs of TTF2, only one showed detectable binding with a very strong binding affinity, which also corresponded to the motif that was predicted to bind to POLE2 according to AF3. No apparent binding can be observed with the NPF motif of ERCC6L.

**Figure 4.**
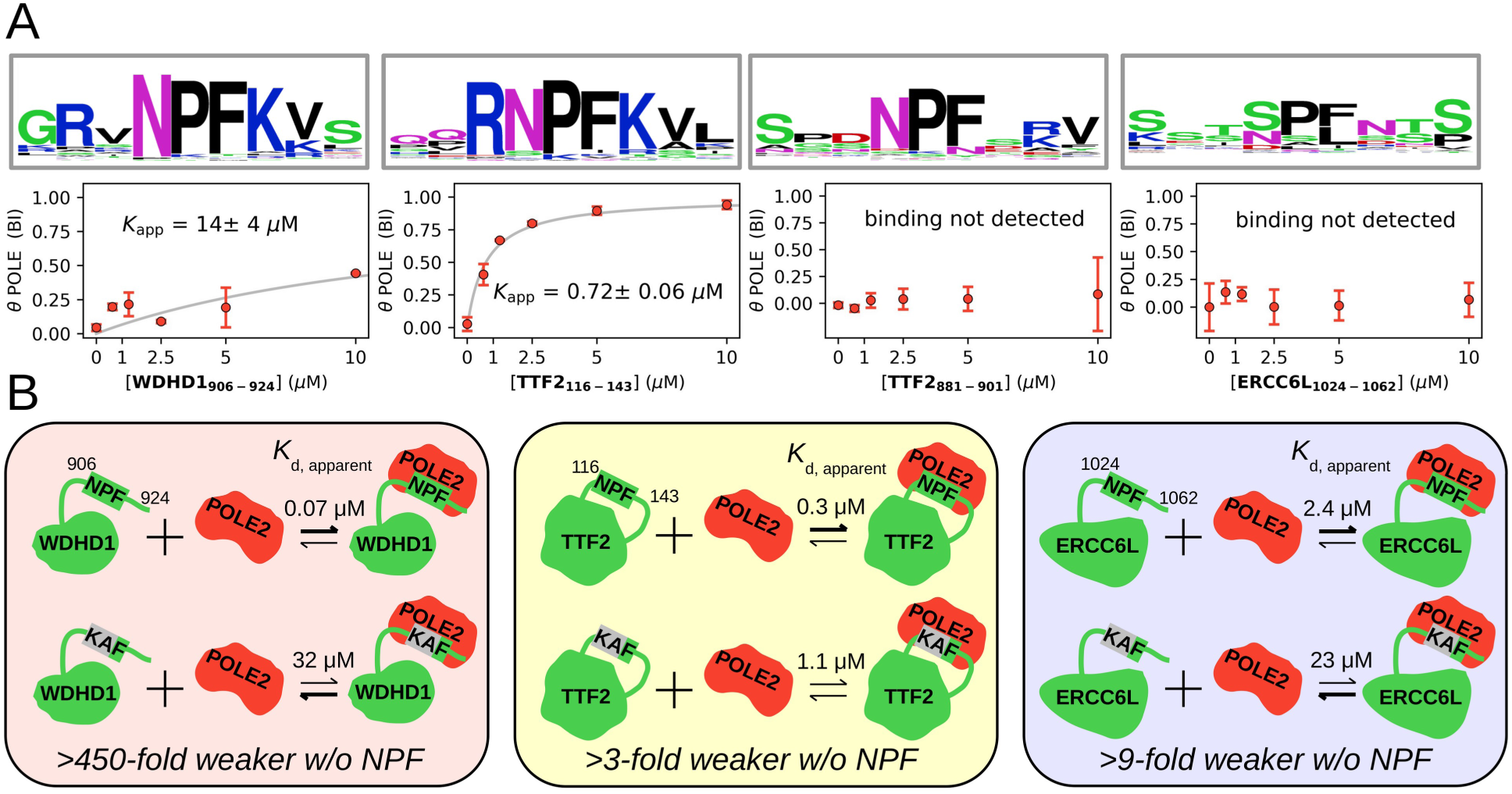
Putative NPF motifs contribute to multiple interactions of POLE2. (A) Four NPF motifs taken from three partners were selected for further studies. Below the evolutionary conservation logo of each, the results of titration nHU experiments are shown. Out of these tested NPF motif peptides, only the one taken from WDHD1 and the first motif of TTF2 shows significant binding to endogenous POLE. (B) Full-length WDHD1, TTF2 and ERCC6L were expressed in HEK293T cells with HA tags with intact or mutated NPF motifs for nHU experiments. In all three cases, mutations of NPF motifs to KAF sequences significantly decreased their binding strength to recombinant POLE2. See Table S2, S3, Figure S8, S12, S13, S14 for more details.

Then, we cloned these three full-length proteins with an HA tag into a mammalian expression vector, produced cell extracts from transfected HEK293T cells and performed nHU titration experiments using purified His_6_-avi-MBP-tagged POLE2 as bait and analyzed the results with dot blot (Figure 4B, S13, S14, S8B, Table S2). The affinities measured in this experiment series were in good agreement with those measured by the single-point proteome scale nHU measurements. Indirect binding of these proteins to POLE2 seems unlikely, as only the target proteins were overexpressed in the cells, without any overexpression of potential intermediate partners. Nevertheless, to validate that these interactions are mediated by the identified NPF motifs, we mutated these NPF sites in the sequences of these proteins into KAF motifs which is expected to abolish their interactions with POLE2 and measured the effect of these mutations with nHU experiments. Only the N-terminal NPF motif of TTF2 was mutated that showed much higher affinity as a synthetic peptide motif. Compared to their wild-type counterparts, the binding of all three mutant proteins were drastically reduced. The largest effect could be measured on WDHD1, which showed a 1000-fold weaker affinity to POLE2 upon mutation. The mutated version of ERCC6L also showed a 10-fold weaker affinity to POLE2. The smallest impact was measured in the case of TTF2, which only showed a 4-fold weaker affinity and could still mediate substantial binding in the presence of the motif-breaking mutation, possibly because of the presence of the secondary weaker NPF motif in its sequence, or because of other interaction regions. Thus, we could validate the presence of functional NPF motifs in three identified partners of POLE2 which bind to the protein directly.

### Intrinsic connection between distinct NPF motif-mediated networks

Substantial evidence was collected to propose an extensive crosstalk between different NPF motif-recognizing proteins (Figure 5A). The NPF motif of BORA could not only interact with AURKA, but also with an EH protein, similarly to WDHD1. POLE2 interacted with a putative NPY motif found in SEC23IP, and this protein is a well-known partner of SEC23/24, which is known to be able to bind to NPF motifs ^19^. The NPF motif of PYGO2 could not only interacted with the core WNT-enhanceosome – SSBP2/4 and LDB1 – but also with EH proteins and AURKA. POLE2 was also found to interact with the WNT-enhanceosome and with many of their known partners, such as SSBP3, LMO4, LEF1, but not with PYGO2. Yet, the most prominent cross-talk was found between EH proteins and POLE2 as our experiments revealed multiple shared partners between these proteins. Particularly, we found that SYNJ1, a known partner of multiple EH proteins, such as EPS15, could also bind to POLE2. This interaction occurs not only through its isolated NPF motif-rich region, but also as a full-length protein. Moreover, other full-length binding partners of EH proteins were found to interact with POLE2 in our screen, including NUMB, EPN2 and AGFG1, all of which are NPF motif-mediated binding partners of EPS15 ^43,44^. Out of these partners, the binding of full-length AGFG1 to POLE2 had been identified in a high-throughput yeast-2-hybrid screen ^45^. It seems that the interaction networks of these unrelated NPF motif recognition proteins intrinsically tend to overlap.

**Figure 5.**
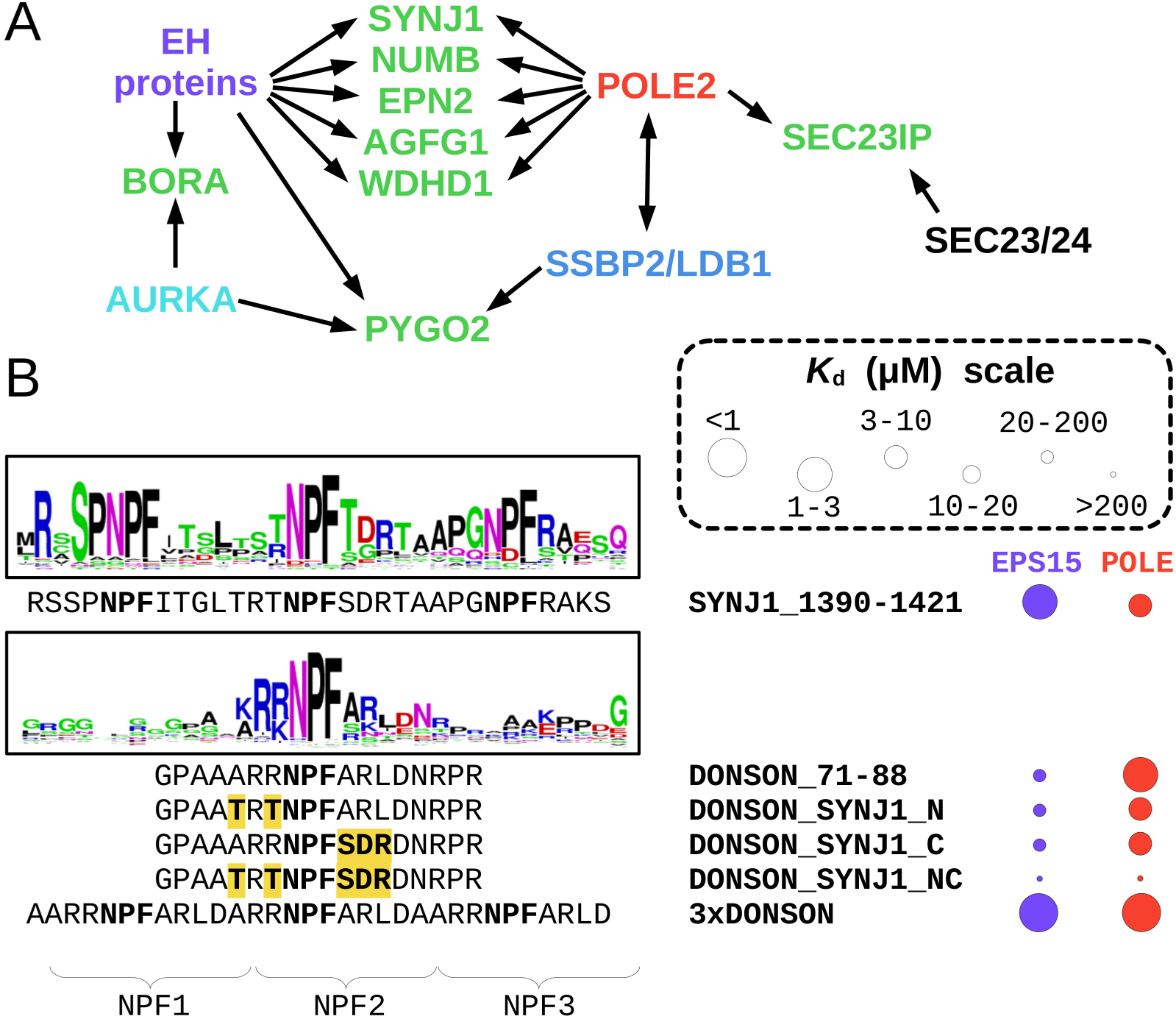
Intrinsic intertwining of discrete NPF networks. (A) Many proteins with NPF motifs were found to interact with multiple unrelated NPF receptors and POLE2 was found to interact with many known NPF motif-mediated partners of other known receptors in the presented experiments. This suggests that instead of multiple discrete cellular NPF networks, a single one exist in cells. (B) To understand the determinants of specificity between EPS15 – an EH protein with three EH domains – and POLE2, a series of DONSON chimera motifs were assayed in nHU experiments against binding to both proteins. This experiment revealed the importance of flanking sequences for efficient POLE2 binding and also the importance of multivalency for EPS15 binding. See Table S2, Figure S15, S16 for more details.

To investigate the determinants of specificity between POLE2 and EPS15, we further studied the NPF motifs of DONSON (that contains a single NPF motif) and SYNJ1 (that contains three NPF motifs) with nHU titration experiments with total Jurkat extract and monitored the binding of endogenous POLE and EPS15 (Figure 5B, S15, S16, Table S2). We found that POLE binds to the DONSON motif with a micromolar *K*_app_ and to the triple motif of SYNJ1 6.5-fold weaker, while EPS15 only interacted with the triple motif of SYNJ1 displaying a comparable micromolar apparent dissociation constant. We tried to turn DONSON into an EPS15 binding partner by replacing the three amino acid long flanking regions of its core NPF motif to the flanking region of the central NPF motif of SYNJ1 either on the N-, or C-terminal, or on both sides. We found that replacing either flanking region decreased the apparent POLE binding affinity by 6-7-fold and when both regions were replaced, the interaction was completely abolished. Thus, POLE2 interaction depends on residues outside the NPF core sequence. However, none of the DONSON-SYNJ1 chimera motifs gained substantial affinity to EPS15, indicating that either the middle NPF motif of SYNJ1 is not recognized by EPS15, or that efficient EPS15 binding requires additional factors.

We hypothesized that this factor may arise from the multivalent nature of EPS15 interactions. To test this, we created a triple NPF motif peptide using the DONSON sequence, following the same spacing as the NPF motifs found in the SYNJ1 sequence. As expected, the 3xDONSON peptide displayed an increased apparent affinity to POLE/POLE2 due to the increased NPF motif density. Unexpectedly, the 3xDONSON peptide also showed a strong gain of affinity to EPS15, even surpassing the natural SYNJ1 peptide motif ligand in binding strength. Multivalency is almost ubiquitously present within the EH protein family, as EH proteins either contain multiple EH domains or form higher-order oligomeric structures, as observed in the case of EHD proteins ^14,15^. Increased apparent affinity of multivalent interaction is a well-known phenomenon and is caused by the increased effective concentration of the interaction sites upon the first binding event ^46^. Still, it is rather surprising that the 3xDONSON displayed even a slightly stronger affinity to EPS15 as one of its endogenous partners, SYNJ1, especially in the light that a single DONSON NPF motif failed to form substantial interaction with EPS15, even when its flanking regions were substituted to one of SYNJ1’s. Thus, it seems that even a suboptimal NPF can efficiently bind to EPS15 once its valency is increased to match EPS15’s. It is also possible that this NPF motif multivalency contributes to some of those seemingly “off-target” interactions observed between POLE2 and conventional EH domain ligands.

### *In vivo* proximity interactome of POLE2

*Ex vivo* holdup experiments are only capable to uncover the intrinsic properties of interaction networks, as the assay uses purified bait molecules and homogeneous cell extracts. Thus, although we could demonstrate that a significant part of the interactomes of various NPF binding proteins can overlap, most of these cross-talks may not occur in their natural cellular environment. To investigate this aspect, we determined the proximity interactome of POLE2 by using BioID experiments in HEK293T cells (Figure 6, Table S4).

**Figure 6.**
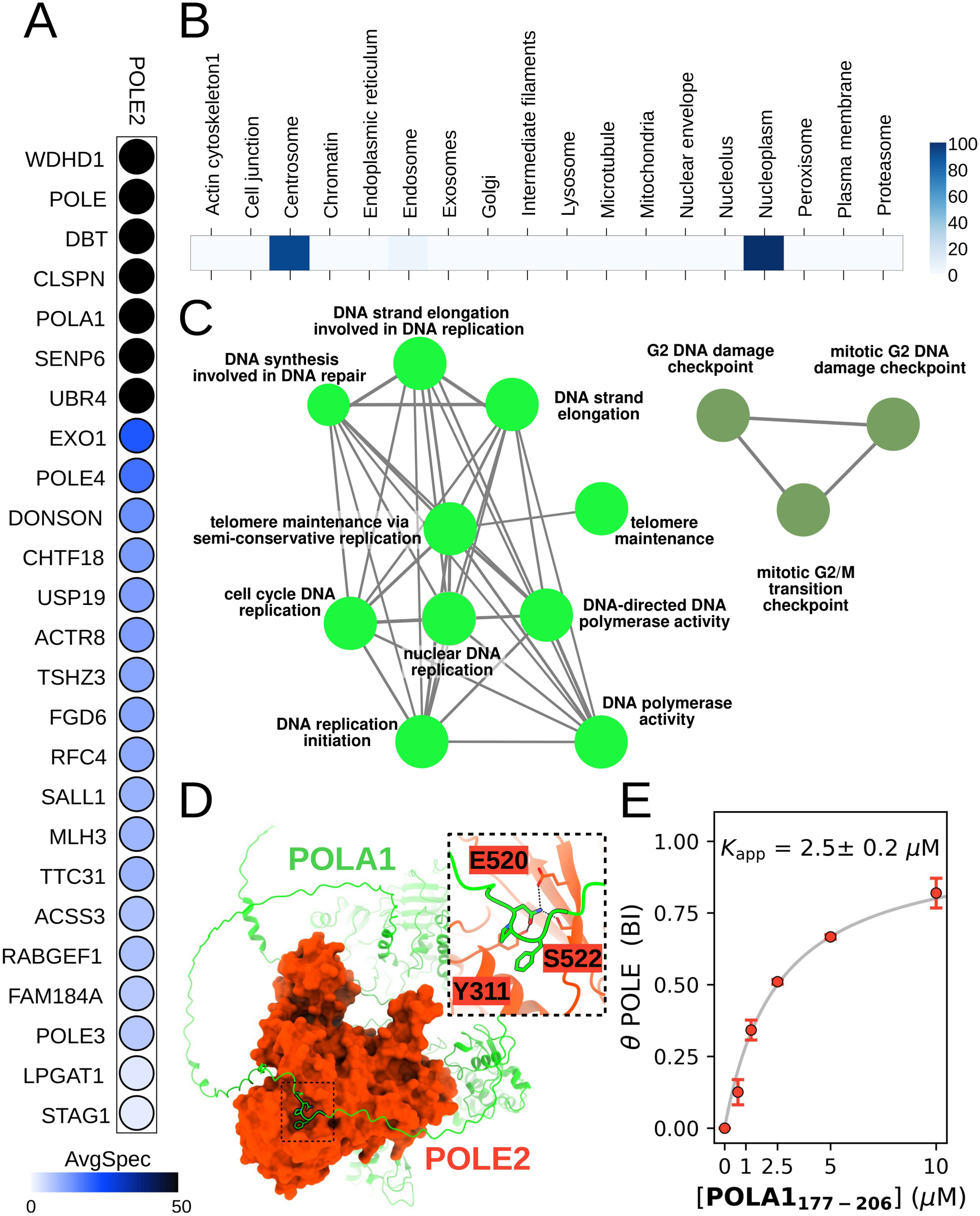
Transient *in vivo* interaction network of POLE2, revealed by BioID. (A) Identified partners of POLE2, ranked by their enrichment. (B) Cellular localization of POLE2 partners reveals that besides nuclear partners, POLE2 also interact with multiple centrosomal proteins, too. (C) Key biological processes mediated by cellular POLE2 partners. Seven partners were identified related to DNA synthesis-related processes and three related to mitotic checkpoint. (D) AlphaFold prediction between POLE2 and POLA1 reveals a potentially NPF-mediated interaction. (E) nHU experiment confirms the interaction between the isolated NPF motif of POLA1 and POLE. See Table S4 for more details.

Using a stringent significance threshold of P < 0.01, the BioID experiment revealed 26 significantly enriched partners in the molecular proximity of POLE2. Only three POLE2 partners were identified by both BioID and nHU: WDHD1, DONSON, and EXO1. Based on the relative enrichment score, the two most enriched POLE2 partners are WDHD1 and POLE, signifying the critical cellular importance of both interactions (Figure 6A). By analyzing the cellular localization of the identified partners, we found that all of them are located either in the nucleoplasm, or in the centrosome, indicating a nuclear and a cytoplasmic POLE2 cellular pool (Figure 6B). We also find a particularly strong enrichment of proteins involved in biological processes related to DNA replication, DNA damage response, and cell cycle control, processes in which POLE has known functions (Figure 6C). Overall, the BioID experiment confirmed that some NPF-motif mediated POLE2 interactions are sufficiently stable to be captured by multiple interactomic approaches. At the same time, BioID also showed that POLE2 is also present in the centrosome, thus we hypothesize that crosstalk between other cytoplasmic NPF receptors and POLE2 may be relevant in certain states of the cell cycle.

While nHU is a powerful approach to identify transient interactions, it has a limited capacity to capture stable complexes, such as the one formed between POLE and POLE2. In contrary, BioID experiments are more suited to detect such interactions and less sensitive to capture the highly transient ones. Consequently, the BioID experiment revealed strong spatial enrichment of POLE, as well as POLE3 and POLE4 in the proximity of POLE2. Beside this ε polymerase complex, only a few core replisome proteins were identified as potential POLE2 partners: WDHD1, Claspin, and POLA1. Out of these, POLA1 is an interaction partner of WDHD1 and may be captured indirectly. However, both Claspin and POLA1 contains a single putative orphan NPF motifs in their disordered protein regions and their direct interaction with POLE2 may be mediated through these sites. In support of this, AF3 successfully docks the putative NPF motif of full-length POLA1 onto the correct binding pocket of POLE2, but it does not find a proper solution in the case of Claspin (Figure 6D). This is in excellent agreement with nHU binding experiments performed with isolated NPF motifs from POLA1 and Claspin that revealed that POLA1 could mediate strong interaction with POLE, while the NPF motif of Claspin could not interact with POLE (Figure 6D, S12, S14, Table S2). In addition to these three replisome proteins, the BioID experiment also identified DONSON as a POLE2 binding partner.

## Discussion

### Motif mediated interactions: distinct networks or coherent interactome

Motif-mediated interactions use limited chemical information space. Their core sequences are short, typically spanning just a few residues where only a few amino acid combinations are permitted. This creates interactomic degeneracy where one motif-binding protein can interact with large numbers of proteins that all contain similar motifs. Certain types of motifs – usually those that are organized around a stable conformation, such as in the case of NPF motifs – are also recognized by not only multiple members of the same motif-binding protein family, but also by multiple unrelated families of different motif-binding proteins. These interaction networks are most often depicted with reductionism as distinct networks with no overlap. However, these networks co-exist in the same cellular environment and they have an intrinsic driving force to merge, because they evolved to capture very similar targets.

In the case of NPF motifs, our study highlights that there is a continuum between the interaction networks of EH proteins, POLE2, and other NPF recognizing protein. Two NPF motifs were found to interact with multiple types of NPF-motif binding proteins out of the five tested. Additionally, we demonstrated that cross-interactions with the EH protein EPS15 can be easily introduced into the NPF motif of DONSON, which naturally displays high selectivity for POLE2. Moreover, the intrinsic interactome of POLE2 revealed its ability to interact with multiple well-known EH protein partners, such as SEC23/24, and even showed interaction with the NPF-binding protein complex SSBP2/LDB1. Although this intrinsic property to form a coherent network clearly exist, we are yet to find evidence that the interaction actually forms in cells. Spatial separation via compartmentalization can prevent the formation of complexes, even those with high intrinsic affinity and in the case of POLE2, we indeed found that the protein exists in a different spatial environment compared to other NPF motif binding proteins. However, under certain circumstances such interactions can still take place and one should not rule out the possibility of the coherent interactome organized around NPF motifs. In support of this, an interaction was already observed between EPS15 and AURKA, two NPF motif-binding proteins, in HEK293T cells ^47^ and between POLE2 and AGFG1, a known EH domain binding protein ^44,45^.

Such interactomic overlap between unrelated motif-binding proteins is not unique for NPF motifs. For instance, we previously described that PDZ-binding motifs can be also bound to kinases, phosphatases and 14-3-3 proteins ^8^, but in principle all motif-mediated interactions that are regulated by post-translational modifications – phosphorylation, acetylation, etc. – display similar interactomic degeneracy. Similarly, many of the known consensus sequences of unrelated motif-binding proteins can overlap, that could, in principle, lead to merged interactomes. For instance, Pro-rich motifs that are often adopt PPII type helical conformations can be recognized by a wide variety of domains, including SH3, WW, EVH1, PTB, or EH domains ^29^. For this reason, we propose to be cautious when relying on consensus sequences to predict interactions of a given motif sequence as these approaches may lead to ambiguous results.

### NPF motifs involved in mechanisms linked to ε DNA polymerase

The ε polymerase holoenzyme is an essential machinery for our cells, most notably responsible for leading strand synthesis during replication (Figure 7A). The complex constitutes of four subunits, the catalytic subunit POLE, the non-catalytic subunit POLE2, as well as two non-essential accessory subunits POLE3 and POLE4 that adopt a histone-like fold. Out of these proteins, most attention was given to the catalytic subunit. This is in part due to the occurrence of multiple clinically observed POLE mutations that lead to Polymerase Proofreading-Associated Polyposis (PPAP), associated with an increased risk of colorectal, stomach, or upper gastrointestinal tract cancers ^48^. However, POLE2 also seems to be critical for proper functioning of the holoenzyme. Upon degradation of POLE in U2OS cells, a decreased POLE2 level was observed and cells were trapped in S phase ^49^. By introducing the C-terminal non-catalytic domain of POLE in these cells, that directly interacts with POLE2, POLE2 levels are not only increased, but replication initiation started with delta polymerase being able to partially replace the function of POLE. This indicates that POLE and POLE2 stabilize each other and that the recruitment of POLE2 at the CDC45-MCM-GINS (CMG) helicase is critical for replication initiation, even in the absence of POLE.

**Figure 7.**
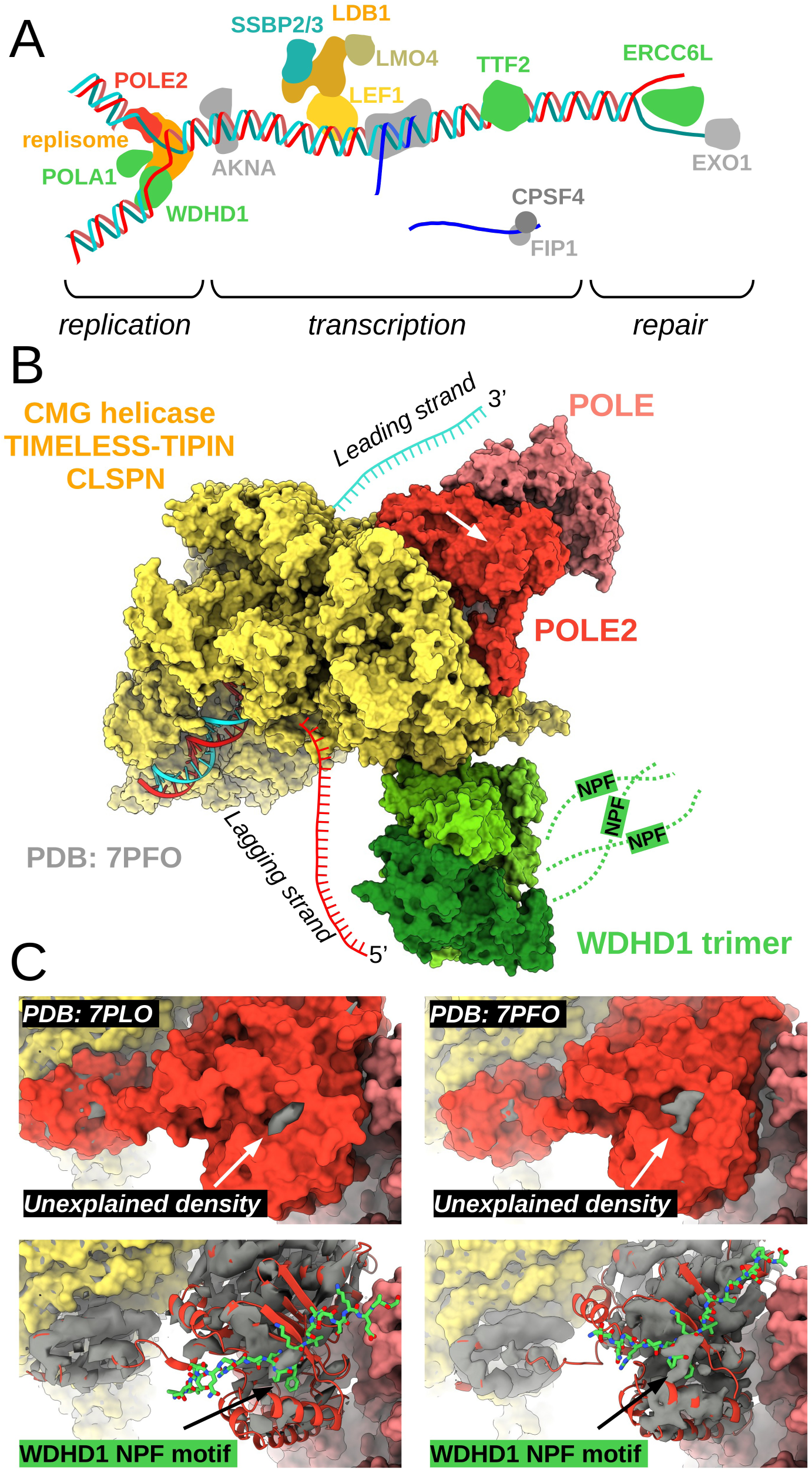
The role of NPF motifs in cellular processes, mediated by POLE2. (A) Identified partners of POLE2 are involved in replication, transcriptional regulation, as well as DNA repair mechanisms. (B) In known replisome structures, the relative position of WDHD1 and POLE2 permits the binding of the WDHD1 NPF motifs. Such stapling would pose a flexible, yet constrained tethering of polymerases responsible for leading and lagging strand synthesis. (C) In the experimental density maps of published POLE2 cryo-EM structures, unexplained densities can be observed on the POLE2 surface perfectly coinciding to the NPF motif sequences docked by AlphaFold.

In the replisome, POLE2 binds the MCM-GINS junction of the CMG helicase and tethers POLE to the machinery ^50^ (Figure 7B). In this conformation, the NPF motif-binding surface of POLE2 is solvent-accessible and in the two human replisome structures both POLE2 and WDHD1 are present. Among all POLE2 partners that contain putative NPF motifs, WDHD1 was the one showing the highest affinity and it also showed the highest enrichment in the proximity labeling experiments. WDHD1 is a trimeric hub protein of the replisome and one of its major functions is to tether alpha DNA polymerase for lagging strand synthesis. Although POLE2 situates on the other end of the replication fork, according to available replisome structures ^50,51^, the NPF motifs in all three copies of the trimeric WDHD1 disordered tails are easily within reach of POLE2. By looking at the experimental low resolution cryo-electron microscopy density maps of the core human replisome (PDB ID 7PFO) or the replisome-CUL2/LRR1 complex (PDB ID 7PLO), unexplained densities can be found on a specific surface of POLE2, but nowhere else in its surrounding region (Figure 7C). This density perfectly coincides with the core NPF motif that was docked on POLE2 by AF3. Thus, it is very likely that in these experimentally captured replisome states, the NPF motif of WDHD1 is bound to POLE2, possibly posing dynamic constraints between the far ends of the replication fork. Similar constraints could be posed through the interactions mediated with POLA1, but this interaction could also play a role at replication initiation through the primase activity of the DNA polymerase alpha complex. In addition to these partners, DONSON is also a critical replisome protein, essential for CMG helicase assembly. At replication initiation, DONSON connects the pre-loading complex – consisting of TOPB1 and GINS1 – to the DNA-bound MCM-CDC45 complex, forming the CMG helicase. Its conserved NPF motif can be important to loosely tether the ε polymerase holoenzyme to forming replisomes ^25,26^. At last, recent studies also identified the NPF motif-dependent interaction between POLE2 and TTF2 and they have found that this interaction is important during mitotic or stalled replisome disassembly by bringing in close proximity a TTF2-bound ubiquitin ligase ^27,28^.

While many of the identified interaction partners of POLE2 that contains NPF motifs are linked to the replisome, the ε polymerase holoenzyme is also known to be involved in other cellular activities, such as nucleotide and base excision repair due to the exonuclease activity of POLE. In alignment with this, most of the interaction partners identified by our holdup experiment are not directly linked to replisome activities. For instance, we identified ERCC6L, a helicase related to ERCC6, which is known to be involved in nucleotide excision repair. In addition, we also identified multiple binding partners of POLE2 that are related to transcriptional control, such as TTF2 with NPF motifs, or SSBP2, a protein without well-conserved NPF motif, but capable of binding NPF motifs. Such interactions may be important during damage-repair, or for replication-transcription control. While these are certainly lesser-known cellular mechanisms of the ε polymerase, NPF motif-mediated interactions may be just as important as those linked to replication.

## Author contribution

SK, KS, AT, TT, and MV designed and performed BioID and all mass-spectrometry measurements and their data analysis. BZ and GG performed all nHU and *in vitro* experiments and analyzed data. NDe and ZD performed bioinformatic analysis. NDa helped in experimental design. MV and GG conceived the project. GG wrote the original draft, and all authors reviewed the manuscript.

## Acknowledgment

We thank the staff of the IGBMC Cell Culture Facility for their help in cell culturing. GG was supported by Inserm, by the ATIP-Avenir program for young group leaders and by the T-ERC STG 2024 (ANR-24-ERCS-0002). This work is part of a project Fondation ARC, funded by Fondation ARC pour la recherche sur le cancer (category PJA1, to GG, 2024). The project was supported by the EKÖP-KDP-24 University Excellence Scholarship Program, the Cooperative Doctoral Program of the Ministry for Culture and Innovation, and the National Research, Development and Innovation Fund (to NoD and ZD).

## Methods

### Cloning

Protein coding sequences (POLE2: UniProt ID P56282, residues 1-527; WDHD1: UniProt ID O75717, residues 1-1129; TTF2: UniProt ID Q9UNY4, residues 1-1162; ERCC6L, UniProt ID Q2NKX8, residues 1-1250) were obtained from cDNA pools using standard protocols. For transient transfection, the protein sequences were cloned into the pCI standard mammalian vector containing a HA-tag for N-terminal tagging with standard restriction cloning. The different mutations were introduced by using standard “quick change” site-directed mutagenesis with Platinum SuperFi II polymerase (Thermo Fisher Scientific). Successful mutagenesis was verified by Sanger-sequencing (Eurofins Genomics Germany GmbH, Ebersberg, Germany). For bacterial expression, POLE2 was cloned as His_6_-AviTag-MBP-TEV-POLE2 construct in a custom bicistronic pET vector that also contains the coding sequence of the E.coli BirA enzyme. The empty His_6_-AviTag-MBP-TEV-STOP vector was used to produce biotinylated MBP for nHU control experiments. For proximity labeling experiments full length POLE2 was cloned to C-terminal (MAC3-tag-C; Addgene Plasmid #185481) destination vector.

### Mammalian cell extract preparation for nHU experiments

Jurkat E6.1 cells (ECACC #88042803, RRID: CVCL_0367) were grown in RPMI (Gibco) medium completed with 10% FBS (Gibco BRL) and 40 μg/ml gentamicin (Gibco/Life Technology). To prepare total cell extracts, Jurkat cells were seeded onto T-175 flasks and grew until 3x10^6^ cells/ml confluency. Cells were collected by 1,000 g x 5 min centrifugation, washed with PBS and then lysed in ice-cold lysis buffer (Hepes-KOH pH 7.5 50 mM, NaCl 150 mM, Triton X-100 1%, cOmplete EDTA-free protease inhibitor cocktail 5x, TCEP 5 mM, glycerol 10%).

HEK293T cells (authenticated with 100% match as ATCC ref. CRL-3216, RRID: CVCL_0063) were grown in Dulbecco’s modified Eagle’s medium (DMEM, Gibco, glucose: 1 g/liter) completed with 10% FCS and 1% Penicillin-Streptomycin (Gibco) on CellBIND flasks (Corning), diluted 1:10 every 3rd/4th day. All cells were kept at 37°C and 5% CO_2_. For transient transfection, cells were seeded on poly-D-lysine (Sigma-Aldrich, ref. P6407) treated 100 mm cell culture dishes (2x10^6^ cells/dish). The next day, the transfection was carried out with JetPRIME reagent (Polyplus) according to the manufacturer’s recommendations using 5 μg DNA and 20 μl of JetPRIME reagent/well. The medium was replaced with fresh medium after 5 hours of treatment with the transfection mixture. Two days after transfection, cells were washed with PBS and then lysed in ice-cold lysis buffer (Hepes-KOH pH 7.5 50 mM, NaCl 150 mM, Triton X-100 1%, cOmplete EDTA-free protease inhibitor cocktail 5x, TCEP 5 mM, glycerol 10%).

Cell suspensions were sonicated 4x20 sec with 1 sec long pulse on ice, then incubated rotating at 4°C for 30 min. The lysates were centrifuged at 12,000 rpm 4°C for 20 min and supernatant was kept. Total protein concentration was measured by standard Bradford method (Bio-Rad Protein Assay Dye Reagent #5000006) using a BSA calibration curve (MP Biomedicals #160069, diluted in lysis buffer) on a Bio-Rad SmartSpec 3000 spectrophotometer instrument. Lysates were diluted to 2 mg/ml concentration, aliquoted and snap-frozen in liquid nitrogen and stored at -80°C until measurement.

### Peptide synthesis

The NPF motif peptide of DONSON (residues 71-88) was chemically synthesized on an ABI 443A synthesizer with standard Fmoc strategy with biotin group attached to the N-terminus via a TTDS (Trioxatridecan-succinamic acid) linker. The NPF motif peptides of SYNJ1 (residues 1390-1421), TTF2 (residues 116-143, or 881-901), WEE1 (residues 228-247), BORA (residues 18-38), ERCC6L (residues 1024-1062), PYGO2 (residues 57-86), and WDHD1 (residues 906-924), as well as all DONSON chimera peptides were obtained commercially from Synpeptide (SYNJ1 peptide) or GenicBio (all other peptides) with an N-terminal biotin label attached to the N-terminus via an Ahx (6-Aminohexanoic Acid) linker. The fluoDONSON peptide (residues 71-88) was N-terminally carboxyfluorescein labeled without using any linker. All peptides were C-terminally amidated and HPLC purified (>95% purity). Predicted peptide mass was confirmed by MS and peptide concentration was determined based on dry weight.

### Protein Expression, Purification

POLE2 was co-expressed with BirA in *E. coli* BL21(DE3) cells. At Isopropyl β-D-1-thiogalactopyranoside (IPTG) induction (0.5 mM IPTG at 18 °C for ON), 50 µM biotin was added to the media. Harvested cells were lysed in a buffer containing 50 mM TRIS pH 7.5, 150-300 mM NaCl, 50 µM biotin, 2 mM 2-mercaptoethanol (BME), cOmplete EDTA-free protease inhibitor cocktail (Roche, Basel, Switzerland), 1% Triton X-100, and trace amount of DNAse, RNAse, and Lysozyme. Lysates were frozen at -20 °C before further purification steps. Lysates were sonicated and centrifuged for clarification. Expressed POLE2 was captured on a HisTrap FF column (Cytiva, 2 x 1 ml), were washed with at least 10 column volume cold wash buffer (50 mM TRIS pH 7.5, 150 mM NaCl, 2 mM BME) before elution with 250 mM imidazole. The Ni-elution was loaded on a pre-equilibrated MBPtrap HP column (Cytiva, 5 ml). The amylose column was washed with 5 column volume cold wash buffer before fractionated elution in a buffer containing 25 mM Hepes pH 7.5, 150 mM NaCl, 1 mM TCEP, 10% glycerol, 5 mM maltose, cOmplete EDTA-free protease inhibitor cocktail. To remove further impurities, POLE2 was ion exchanged with a Q HP column (Cytiva) using a linear gradient from buffer A (50 mM Tris pH7.5, 25 mM NaCl, 2 mM BME) to buffer B (50 mM Tris pH7.5, 1 M NaCl, 2 mM BME) with full-length POLE2 eluting in a single peak at ∼20% buffer B. Biotinylated MBP was expressed similarly, but was only purified by the two affinity chromatography steps. 10% glycerol, 5 mM TCEP was supplemented to the eluted proteins. The concentration of proteins was determined by their UV absorption at 280 nm before aliquots were snap frozen in liquid nitrogen and were stored at -80 °C.

### Single-point and titration nHU experiments

Single-point nHU experiments with peptide baits were carried out at ∼10 μM estimated bait concentration. For this, 50 μl streptavidin resin (Streptavidin Sepharose High Performance, Cytiva) was incubated with 250 μl 70 μM biotinylated peptides (or biotin for controls) in 0.5 ml Protein LoBind tubes (Eppendorf). After mixing, reaction mixtures were incubated overnight at 4 °C, or for 1 hour at room temperature. After the incubation, all resins were washed with 9 resin volume (450 μl) holdup buffer (50 mM Tris pH 7.5, 300 mM NaCl, 1 mM TCEP, .22 filtered). The washed resins were then mixed with 25 μl 1 mM biotin solution, diluted in 10 resin volume holdup buffer and were incubated for 5 minutes at room temperature. Then, the resulting resins were washed four times in total with 9 resin volume holdup buffer. The resulting bait-saturated resins were mixed with 200 μl 2 mg/ml Jurkat extracts and were incubated at 4 °C for 2 h with constant mild agitation. After the incubation ended, the solid and liquid phases were separated by a brief centrifugation (15 sec, 2000 g) and 130 μl of the supernatant was recovered rapidly. Then, to minimize carryover contamination from resin, the recovered supernatants were centrifuged one more time and 100 μl of the supernatant was recovered that was subjected for mass spectrometry analysis.

Single-point nHU experiments with protein baits were carried out nearly identically with two key modifications. First, the resin saturation step was carried out by mixing 50 μl streptavidin resin with 0.5 - 1 ml 50 μM biotinylated protein solution and the mixture was incubated for 1.5 hours at 4 °C with constant agitation. Second, instead of using estimated bait concentrations, we measured it experimentally. An extra aliquot of resin stock was prepared alongside the nHU samples that were identically processed that were used to eluate the immobilized proteins under harsh conditions, in order to experimentally quantify the *C*_bait_ by using densitometry of Coomassie stained SDS-PAGE that included a protein calibration series. For this, 50 μl bait-saturated resin aliquots were mixed with 200 μl 4x Laemmli Sample Buffer (120 mM Tris-HCl pH 7, 8% SDS, 100 mM DTT, 32% glycerol, 0.004% bromphenol blue, 1% β-mercaptoethanol), instead of cell extract. After 15 min incubation at 95 °C, these samples were analyzed by SDS-PAGE using protein calibration standard, made from recombinant MBP. Densitometry was carried out after Coomassie staining on raw Tif images by using Fiji ImageJ 1.53c and experimental bait concentration were determined by adjusting for molecular weight differences between MBP and MBP-POLE2 constructs. This experiment revealed that the measured bait concentration (26-40 μM) is slightly higher than the empirically estimated (10 μM).

Titration holdup experiments were carried out as described above using 25 μl saturated resins prepared ^35^. Briefly, we mixed control, or bait-saturated resins at various proportions by keeping the total resin amount and resin-analyte ratio constant during the experiment (for 25 μl we used 100 μl cell extract as analyte). In titration experiments with protein baits, bait concentration was determined as described above for the saturated resin stocks. Experiments were carried out at 4 °C for 2 h and recovered supernatants were subjected to Western blot or Dot Blot analysis.

### Liquid Chromatography–Mass Spectrometry analysis of nHU experiments

The 100 µl nHU samples were TCA precipitated by adding 20 µl 100% TCA (#100807, Merck). Immediately after TCA addition the samples were mixed well with vortex and left overnight to 4°C to precipitate. The following day the samples were centrifuged at 13 000 rpm for 15 min at 4°C. The protein pellets were washed with 400 µl ice-cold 100% acetone, and centrifuged 13 000 rpm for 8 min at 4°C. The supernant was discarded and the pellets washed again with 400 µl ice-cold 100% acetone followed by centrifugation at 13 000 rpm for 5 min at 4°C. After centrifugation the supernanant was carefully removed by pipetting and pellets were air dried. To drive off acetone the pellets were placed in 95°C heat block for 5-10 min until no liquid was visible. 50 µl 8M Urea was added to the pellets, mixed with vortexing and put to -20°C to solubilize. The following day the samples were vortexed and left overnight to 4°C to solubilize. After the incubation 8M Urea was diluted with 100 mM Tris-HCl, pH 8.0 to 4M by adding 50 µl buffer, mixing by vortexing. Proteins were reduced with 5 mM (11 µl of 50 mM stock) TCEP for 30 min at 37°C, and alkylated with 10 mM ioodoacetamide (12 µl of 100 mM stock) for 30 min in the dark at room temperature. Urea concentration was diluted to ∼1 M by adding 300 µl 100 mM Tris-HCl, pH 8.0. The pH of the samples was controlled and 1 µg Trypsin/Lys-C Mix (V5073, c=0.5 µg/µl, Promega) was added to each sample and mixed by vortexing. Samples were digested overnight at 37°C. Peptide mixtures were then desalted on a C18 mini columns (#HUM S18V, Higgins Analytical, Inc.), and dried in a centrifuge concentrator (Concentrator Plus, Eppendorf).

For MS acquisition the dried peptides were resuspended in 40 µl Buffer A (0.1% (vol/vol) TFA, 1% (vol/vol) acetonitrile (#83640.320, VWR) in HPLC grade water (#10505904, Fisher Scientific). For the DIA analysis, the resuspended peptides were further diluted 1:20 in buffer A1 (1% formic acid in HPLC water). 20 µl was loaded into an EvotipPure (Evosep, Denmark) according to the manufacturer’s instructions and run at the same time as three technical replicates.

The samples were analyzed using the Evosep One liquid chromatography system coupled to a hybrid trapped ion mobility quadrupole TOF mass spectrometer (Bruker timsTOF Pro2, Bruker Daltonics) ^52^ via a CaptiveSpray nano-electrospray ion source (Bruker Daltonics). An 8 cm × 150 µm column with 1.5 µm C18 beads (EV1109, Evosep) was used for peptide separation with the 60 samples per day method (21 min gradient time). Mobile phases A and B were 0.1 % formic acid in water and 0.1 % formic acid in acetonitrile, respectively. The MS analysis was performed in the positive-ion mode with dia-PASEF method ^53^ with sample optimized data independent analysis (dia) scan parameters. We performed DDA in PASEF mode from a pooled sample to be able to adjust dia-PASEF parameters optimally. To perform sample-specific dia-PASEF parameter adjustment, the default dia-short-gradient acquisition methods were adjusted based on the sample-specific DDA-PASEF run with the software “tims Control” (Bruker Daltonics). The ion mobility windows were set to best match the ion cloud density from the sample type-specific DDA-runs. The following parameters were used in dia-PASEF runs: TIMS settings 1/KO 0.85-1.30, Ramp time 100 ms, Accumulation time 100 ms, Duty cycle 100%, and dia-PASEF settings cycle time estimate 1.17 s, MS1 ramps 1, MS/MS ramps 10, MS/MS windows 18, mass range 357.2-1257.2 Da, Mobility range 0.85-1.30 1/K0. For collision energy at 0.60 1/KO 20 eV and at 1.60 1/KO 59 eV were used.

To analyze diaPASEF data, the raw data (.d) were processed with DIA-NN v1.8.1 ^54,55^ utilizing spectral library generated from the UniProt human proteome (UP000005640, downloaded 05.04.2022 as a FASTA file, 20358 proteins). During library generation, the following settings were used with fixed modifications: carbamidomethyl (C); variable modifications: acetyl (protein N-term), oxidation (M); enzyme:Trypsin/P; maximum missed cleavages: 1; mass accuracy fixed to 1.5e-05 (MS2) and 1.5e-05 (MS1); Fragment m/z set to 100-1700; peptide length set to 7-30; precursor m/z set to 300-1800; Pre cursor changes set to 2-4; protein inference not performed. All other settings were left to default.

### Statistical and bioinformatics analysis of the proteomics data for nHU samples

The input file to further dia data analysis was the DIA-NN Report.pg_matrix. The measured protein intensities were normalized based on median intensities of the entire dataset to correct minor loading differences. For statistical tests and enrichment calculations, not detectable intensity values were treated with an imputation method, where the missing values were replaced by random values similar to the 10% of the lowest intensity values present in the entire dataset. Unpaired two tailed T-test, assuming equal variance, were performed on obtained log_2_ intensities. Obtained fold-change values were converted to apparent affinities using the hyperbolic binding equation (Equation 2) for those interactions that were found to be significant above (2 σ) thresholds as described before ^33,35^.

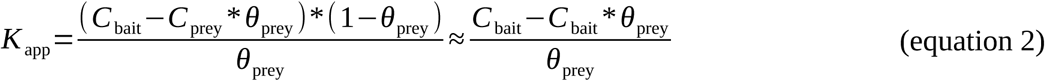

Note that interactions below this stringent significance threshold may be still biologically important and the thresholds were set to minimize type 1 error (false positive). Interactions with weak affinities will result in small depletion values and interaction partners that are difficult to quantify will lead to low statistical significance, both resulting in type 2 error (false negative). Imputation is also a clear source of type 2 error. For example, endogenous DONSON did not showed as a significant partner of POLE2. However, it is clear at closer inspection that this is due to the difficulty of DONSON quantification by mass spectrometry in our nHU samples and due to an imputation step of the MS analysis. In this case, only two low abundance peptides of DONSON were detected. Out of these, one peptide was completely absent in the POLE2-depleted nHU samples and the other displayed an approximately 22% depletion, however this value is based on a single quantification. Nevertheless, this depletion equals to a 92 μM dissociation constant, that is an affinity comparable to the other NPF motif-mediated partners of MBP-POLE2.

### Estimating affinities of indirectly captured complexes

In all nHU-MS experiments with NPF motif peptides where POLE/POLE2 binding was detected, the significance level of POLE depletion was higher. This is possibly due to the larger size of POLE compared to POLE2, causing the easier detection of POLE and more robust quantification by MS. On one occasion, this resulted in POLE2 being measured as significantly depleted, but the depletion fell below our statistical significance threshold. Regardless, in all experiments done with DONSON, WDHD1, or SYNJ1 the degree of binding of POLE2 was found to be consistently higher than of POLE. This information itself strongly implies that the NPF motif interacts directly with POLE2 and that this binding can occur regardless whether POLE2 is bound to POLE, or not. Moreover, this shows that the POLE-POLE2 heterodimer is not an obligate dimer and display some dynamic properties. As POLE shows smaller depletion, it also displays weaker apparent affinity against the same NPF motif baits compared to POLE2. If we assume that POLE binds to POLE2 independently to the POLE2-NPF motif interaction, the measured differences in the degree of binding for POLE and POLE2 can even help us to estimate the apparent binding strength of the POLE-POLE2 interaction, according to

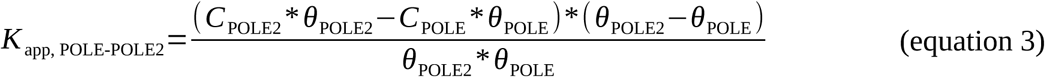

where *C*_POLE2_ and *C*_POLE_ are the total POLE2 and POLE concentrations in the extract, and *θ*_POLE2_ and *θ*_POLE_ are the measured depletion values of the two proteins against the same bait. Based on proteomic studies, POLE and POLE2 are expressed in stoichiometric amounts in cells and they form a heterodimer, approximately at 100 nM cellular concentration ^56^. However, our cell extracts can be considered as 100-fold diluted cells resulting an approximately 1 nM concentration for POLE and POLE2. Given this, as well as the MS-based prey depletion measurements, equation 1 gives an estimated dissociation constant of the POLE2-POLE interaction between 0.1 and 0.3 nM, calculated for the different NPF motif baits. As of note, we could identify multiple complexes as binding partners of NPF motifs, or POLE2 by comparing lists of interaction partners with interactomic datbases ^45,47,57,56^ (Figure S9). It would be theoretically possible to estimate affinities for interactions within these complexes in a similar fashion.

### Western and Dot blot experiments

For Western blot, samples were mixed with 4x Laemmli Sample Buffer in 3:1 ratio. Equal amounts of samples were loaded on 6-10% acrylamide-gels (generally 16 μg). Transfer was done into PVDF membranes using a Trans-Blot Turbo Transfer System and Trans-Blot Turbo RTA Transfer Kit (BioRad, #1704275). For Dot blot, nHU samples were directly loaded on Nitrocellulose membranes using 48 Sample vacuum manifold (Cleaver Scientific) following the manufacturer’s recommendations.

After 1 hour of blocking in 5% milk, membranes were incubated overnight at 4°C with primary antibodies in 5% milk-TBS-Tween. The following antibodies and dilutions were used: POLE antibody (PA5-78113, Thermo Fisher Scientific, RRID: AB_2736449, dilution: 1:2,000), EPS15 antibody (D3K8R, #12460, Cell Signalling Technologies, RRID: AB_2797926, dilution: 1:1,000), HA antibody (HA.11 epitope tag, BioLegend, ref. 901501, RRID: AB_2565006, dilution 1:5,000). Then, the membranes were washed three times with TBS-Tween and incubated at RT for 1 h in secondary antibody (goat anti-rabbit(H+L) #111-035-003 RRID: AB_2313567 or goat antimouse(H+L) #115-035-146 RRID: AB_2307392) in 5% milk-TBS-Tween (dilution: 1:10,000). After washing three times with TBS-Tween, membranes were exposed to chemiluminescent HRP substrate (Immobilon, #WBKLS0100) and captured in docking system (Amersham Imager 600 or 800, GE). When membranes were re-exposed to other primary antibody raised in the same species, the membranes were stripped in 30 min with mild stripping buffer [glycine (15 g/liter), SDS (1 g/liter), and 1% Tween 20 (pH 2.2)] to remove primary signal. When membranes were re-exposed to a primary antibody raised in a different species, membranes were treated with 15% H_2_O_2_ for 30 seconds to remove secondary signal. After a 30 min re-blocking in 5% milk-TBS-Tween, membranes were incubated overnight with the following primary antibody and continued the blot as described above. For control, anti-α-tubulin primary antibody (Sigma-Aldrich, ref. T9026, RRID: AB_477593, dilution: 1:5,000) or anti-GAPDH antibody (clone 6C5, MAB374, Sigma-Aldrich, RRID: AB_2107445, 1:5000) were used (in 5% milk-TBS-Tween for 1 h).

Densitometry was carried out on raw Tif images by using Fiji ImageJ 1.53c. Data fitting of titration experiments were performed in QtiPlot and data visualization was done using Matplotlib. For interactions following partial binding, the following formula was used:

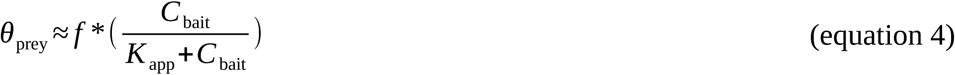

where the *f* factor is the “fraction of binding capable prey”. (For interactions displaying complete binding, the f value was set to 1, which is identical to the equation shown on Figure 2A.)

### Direct and competitive Fluorescence Polarization experiments

Fluorescence polarization (FP) was measured with a PHERAstar microplate reader by using 485 ± 20 nm and 528 ± 20 nm band-pass filters for excitation and emission, respectively. In direct FP measurements, a dilution series of MBP-POLE2 was prepared in 96 well plates (96 well skirted PCR plate, 4ti-0740, 4titude, Wotton, UK) in a 20 mM HEPES pH 7.5 buffer that contained 150 mM NaCl, 0.5 mM TCEP, 0.01% Tween 20 and 50 nM fluoDONSON peptide. The volume of the dilution series was 40 μl, which was later divided into three technical replicates of 10 μl during transferring to 384 well micro-plates (low binding microplate, 384 well, E18063G5, Greiner Bio-One, Kremsmünster, Austria). In total, the polarization of the probe was measured at eight different protein concentrations (whereas one contained no protein and corresponded to the free peptide). In competitive FP measurements, the same buffer was supplemented with 3.85 μM POLE2 protein to achieve sufficient complex formation to be able to quantify competition event. Then, this mixture was used for preparing a dilution series of the competitor not-fluorescent DONSON peptide and the measurement was carried out identically as in the direct experiment. Analysis of FP experiments were carried out using ProFit, an in-house developed, Python-based fitting program ^58^. The dissociation constants of the direct and competitive FP experiments were obtained by fitting the measured direct data with a quadratic binding equation first and by fitting the measured competitive data with a competitive equation, using several obtained parameters from the first fit.

Note that unlike titration holdup experiments, conventional fluorescence polarization experiments are unsuitable for detecting partial binding, because this assay only measures the state of the fluorescent molecules.

### Motif identification and analysis

To find all putative NPF motifs in the proteome, the SLiMSearch 4 program was used using default parameters and the NPF consensus sequence with 5 flanking residues on each end ^37^. The obtained list was cross checked with the list of significant POLE2 interaction partners based on UniProt identifiers to identify all partners with putative motifs. To visualize evolutionary conservation of selected NPF motifs, automatic sequence alignments of orthologous sequences were performed using ProViz using metzoan sequences and the human sequence template and sequence logos were created using WebLogo ^59^.

The sequences were collected for the POLE2 interactome from UniProt ^60^, and disordered regions were identified using the following criteria: positions classified as disordered were retained if they did not overlap with predicted Pfam ^61^ domains and they overlapped with the experimentally verified disordered region based on MobiDB ^62^ Otherwise, regions predicted as disordered by AlphaFold2-rsa (threshold > 0.582) or when AlphaFold2 ^63^ structures were unavailable, based on IUPred2A ^64^ scores (threshold > 0.4) were considered as disordered. Regions with less than 5 amino acids were omitted.

Motif discovery was performed on disordered regions using the STREME tool ^40^ with the following parameters: a minimum motif length of 3, a maximum length of 15, and a maximum of 25 motifs to be identified. Vertebrata conservation data were calculated as described previously and were retrieved from the DisCanVis web server ^65^. For each predicted motif, the conservation scores were averaged of their corresponding sites and the predicted motifs were reordered based on the mean conservation score.

### AlphaFold predictions

Structure predictions of binary complexes were performed using the AlphaFold 3 online server with default settings ^36^. To calculate the complexes of peptide-bound POLE2 structures, the sequence of full-length POLE2 was used together with the NPF motif peptide sequences. To calculate the structures of binary complexes of full-length proteins, the sequences of POLE2 was used with each interaction partners. All obtained structures were analyzed by evaluating if the NPF motif-binding pocket of POLE2 was occupied by the interaction partner. In all instances when this pocket was utilized for interactions, the pLDDT scores of the docked NPF or NPY motifs were outstanding compared to their environment and were similar to the scores of the globular POLE2 regions. The obtained peptide-bound structure of POLE2-WDHD1 was superposed to the C-terminal domain POLE2 of the cryo-EM structures and no further adjustment was made to improve the fit of the superposed NPF motif peptide segment into the experimental density maps.

As of note, a possible explanation for why AlphaFold was able to provide a high confidence structure prediction is because a pseudo NPF motif-mediated interaction of POLE2 was already observed as a crystallographic artifact. A crystal structure of POLE2 captured a crystallographic dimer where the C-terminal QGF motif of POLE2 is bound to the NPF binding pocket of the non-crystallographic symmetry-related POLE2 molecule ^66^ (Figure S17). In this structure, the QGF motif similarly forms a turn conformation, where main-chain of Q525 is jointly coordinated by E520 and S522 from the other molecule and where F527 is bound to the same hydrophobic pocket formed by Y311.

### BioID experiments

To generate tetracycline-inducible stable cell line expressing C-terminally MAC3-tagged POLE2 and GFP control were introduced into Flp-In T-REx 293 cells (Life Technologies, Carlsbad, CA) ^67^. Approximately 1 × 10^7^, per replicate, Flp-In T-REx 293 cells stably expressing MAC3C-tagged POLE2 were induced with 2 μg/ml tetracycline for 24 h to induce the transgene expression, and 50 μM biotin added to 3 h before the harvesting to induce the biotinylation. The cells were washed three times with 1 x PBS and harvested with 1 mM EDTA in 1× PBS. The harvested cells were pelleted using centrifugation, snap frozen in liquid nitrogen, and stored at −80 °C. The samples were then suspended in 1.5 ml of lysis buffer (0.5% IGEPAL, 50 mM HEPES, pH 8.0, 150 mM NaCl, 50 mM NaF, 1.5 mM NaVO_3_, 5 mM EDTA, 0.1% SDS, 0.5 mM PMSF, protease inhibitors, and 80 U/ml benzonase nuclease (Santa Cruz Biotechnology)) on ice. Lysis was followed by incubation on ice for 15 min and three cycles of sonication (3 min) and incubation (5 min) on ice. The samples were cleared by centrifugation, and the supernatants were poured into microspin columns (Bio-Rad) that were preloaded with 250 μl of (50 % slurry) Strep-Tactin beads (IBA GmbH) and allowed to drain under gravity. The beads were washed 3 times with 1 ml lysis buffer and then 4 times with 1 ml HENN (50 mM Hepes pH 8.0, 5 mM EDTA, 150 mM NaCl, 50 mM NaF) buffer. The purified proteins were eluted from the beads with 700 μl of wash buffer containing 0.4 mM biotin. To reduce and alkylate the cysteine bonds, the proteins were treated with a final concentration of 5 mM TCEP (tris(2-carboxyethyl) phosphine) 20 min at 37 °C by shaking and 10 mM IAA (iodoacetamide) for 20 min in dark. Finally, the proteins were digested into tryptic peptides by incubation with 1 μg sequencing grade trypsin (Promega, V5113) overnight at 37 °C by shaking. The digested peptides were purified using C-18 microspin columns (Higgins Analytical, Inc.) as instructed by the manufacturer. For the mass spectrometry analysis, the vacuum-dried samples were dissolved in buffer A (1% acetonitrile and 0.1% trifluoroacetic acid in MS-grade water).

### Liquid Chromatography–Mass Spectrometry analysis of BioID experiments

The Evosep One liquid chromatography system was utilized to analyze desalted peptide samples. It was coupled to a hybrid trapped ion mobility quadrupole TOF mass spectrometer (Bruker timsTOF Pro 2) via a CaptiveSpray nano-electrospray ion source. Peptides were separated using an 8 cm × 150 μm column packed with 1.5 μm C18 beads (EV1109, Evosep) utilizing the 60-SPD (sample-per-day) method with a 21-min gradient time. Mobile phases A and B were prepared as 0.1% formic acid in water and 0.1% formic acid in acetonitrile, respectively. For all BioID samples, mass spectrometry (MS) analysis was performed in positive-ion mode using a data-dependent acquisition (DDA) strategy in PASEF (Parallel Accumulation Serial Fragmentation) mode, employing the DDA-PASEF-short_gradient_0.5s-cycletime method. The raw data files were processed utilizing FragPipe v22.0 in DDA+ mode with MSFragger-4.1 ^68^ using human protein fasta files containing 40,970 entries (20,485 decoys: 50%) from the UniProtKB database (downloaded on October 21, 2024). Carbamidomethylation of cysteine residues was used as static modification while amino-terminal acetylation and oxidation of methionine were used as the dynamic modification. Biotinylation of lysine and N-termini were set as variable modifications. Trypsin was selected as the enzyme with allowance for up to two missed cleavages. The instrument and label-free quantification parameters remained at their default settings. Lastly, the results output consisted of PSM values derived from peptides with a false discovery rate (FDR) below 0.01 as gener ated by Philosopher.

### Identification of High-Confidence Protein-Protein Interactions of BioID experiments

In-house python script, incorporated with Significance Analysis of INTeractome (SAINT) -express version 3.6.3 ^69^ was used as a statistical approach for identification of specific high-confidence interactions from BioID. High-confidence interactions (HCIs) were defined by an estimated protein-level Bayesian FDR (BFDR) of ≤0.01. Interactions that passed the filter with any bait were rescued for other baits as well. Furthermore, we used our own in-house contaminant GFP library and CRAPome database (version 2.0) ^70^. Preys detected in over 20% of the GFP library or CRAPome samples were deemed contaminants and removed unless their spectral counts in the sample runs were over 3 times higher than the library average. Preys with an average spectral count of less than three were also removed.

## Data Availability

All data needed to evaluate the conclusions in the paper are present in the paper, in the Supplementary Materials and in public databases. All raw LC-MS/MS data have been deposited to the ProteomeXchange via the MassIVE database under the ID of MSV000097307.

## Supplemental Figures

**Figure S1.**
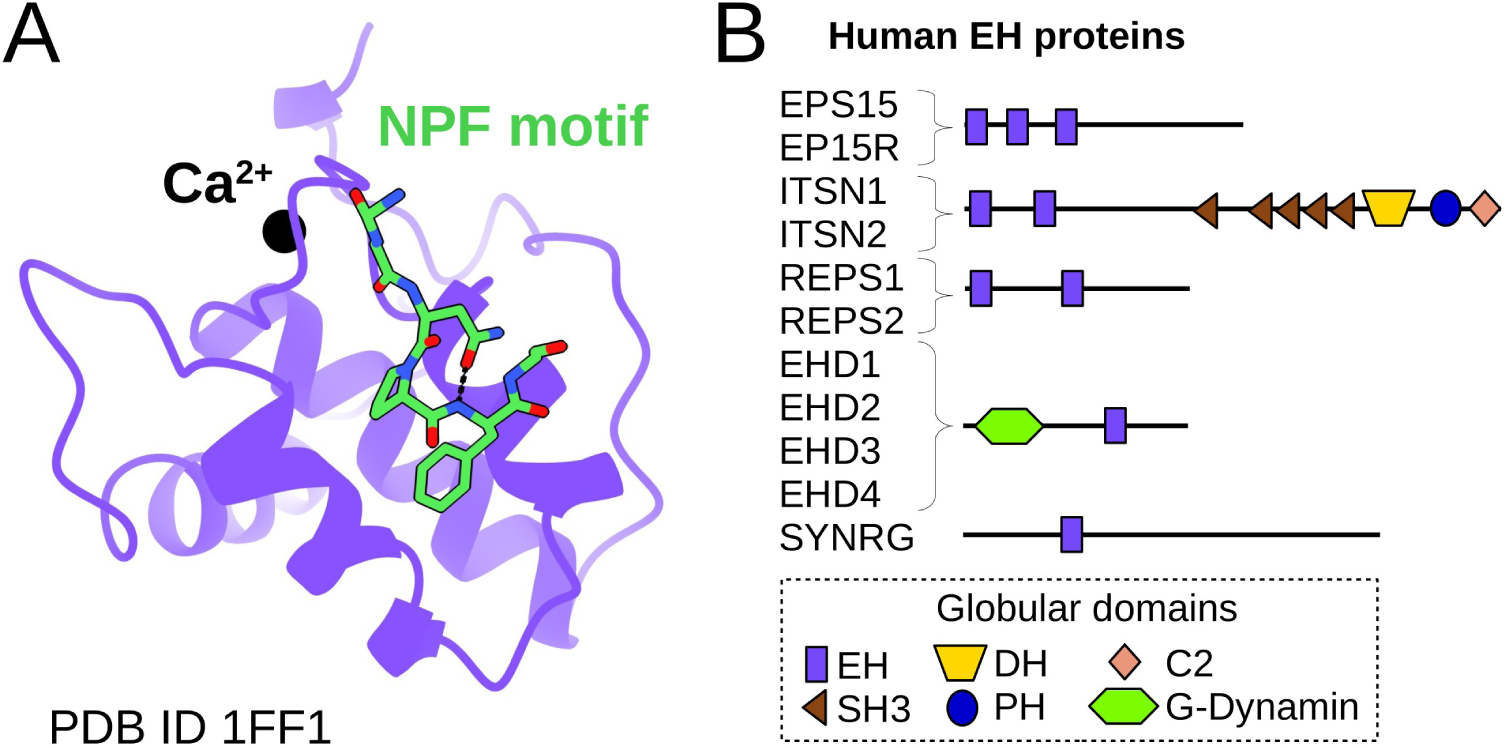
EH proteins and NPF motif binding. (A) Structural snapshot of an interaction formed between an EH domain and an NPF motif. (B) Domain architecture of the human EH protein family.

**Figure S2.**
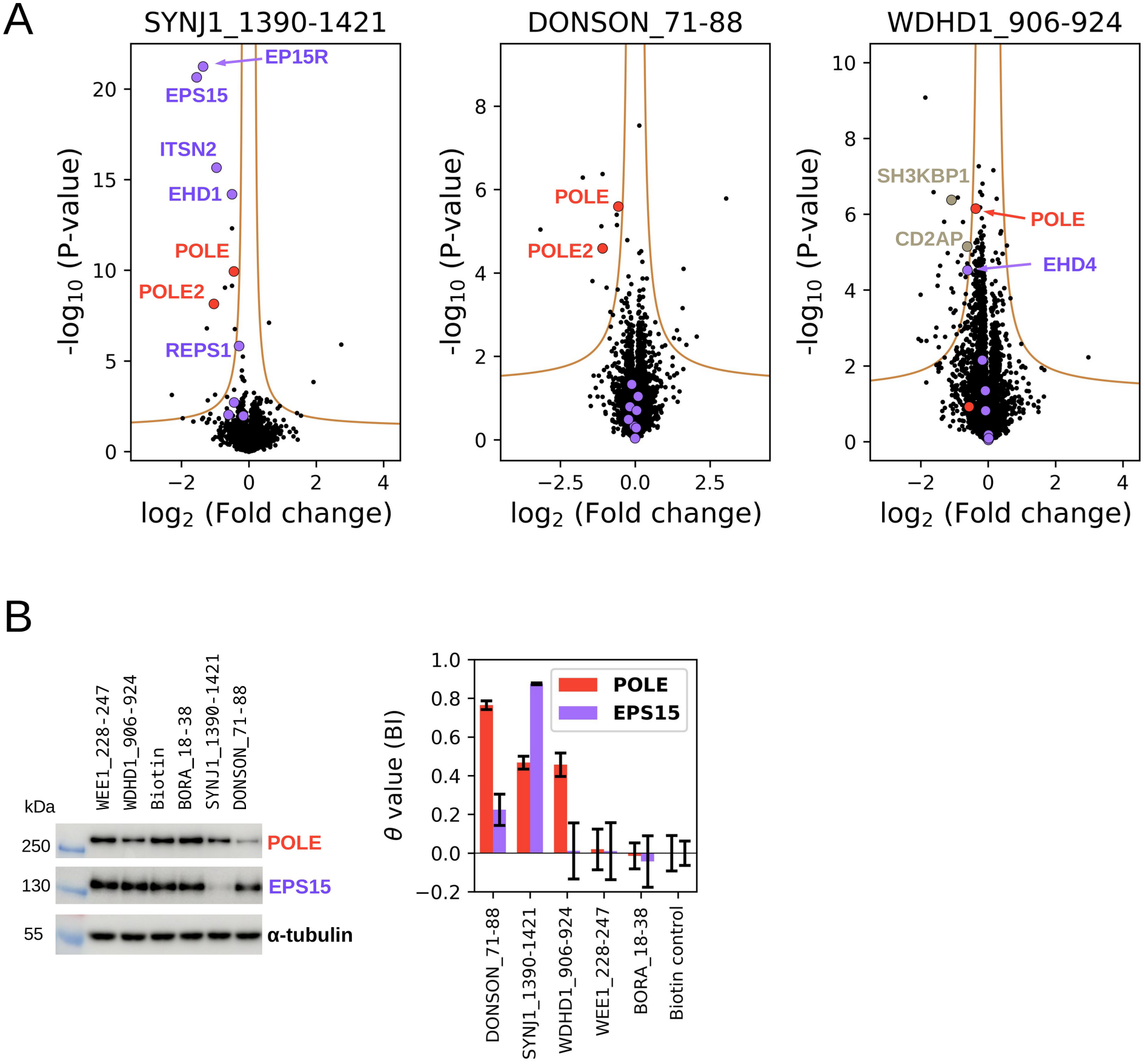
Complementary nHU experiments confirm the interaction between multiple NPF motifs and POLE/POLE2. (A) Independent nHU-MS measurements carried out with three NPF motif peptide baits. Experiments were carried out with 2 biological and 3 technical replicates (total N = 6). (B) Summary of validation experiments analyzed by Western blot. The binding of endogenous POLE and EPS15 is monitored for five selected NPF motifs at a fixed bait concentration. The binding of POLE is clearly detected in the case of bait peptides taken from DONSON, WDHD1, and SYNJ1, wile EPS15 could only interact with SYNJ1 and to a lesser degree DONSON. Western blots were performed in technical duplicates.

**Figure S3.**
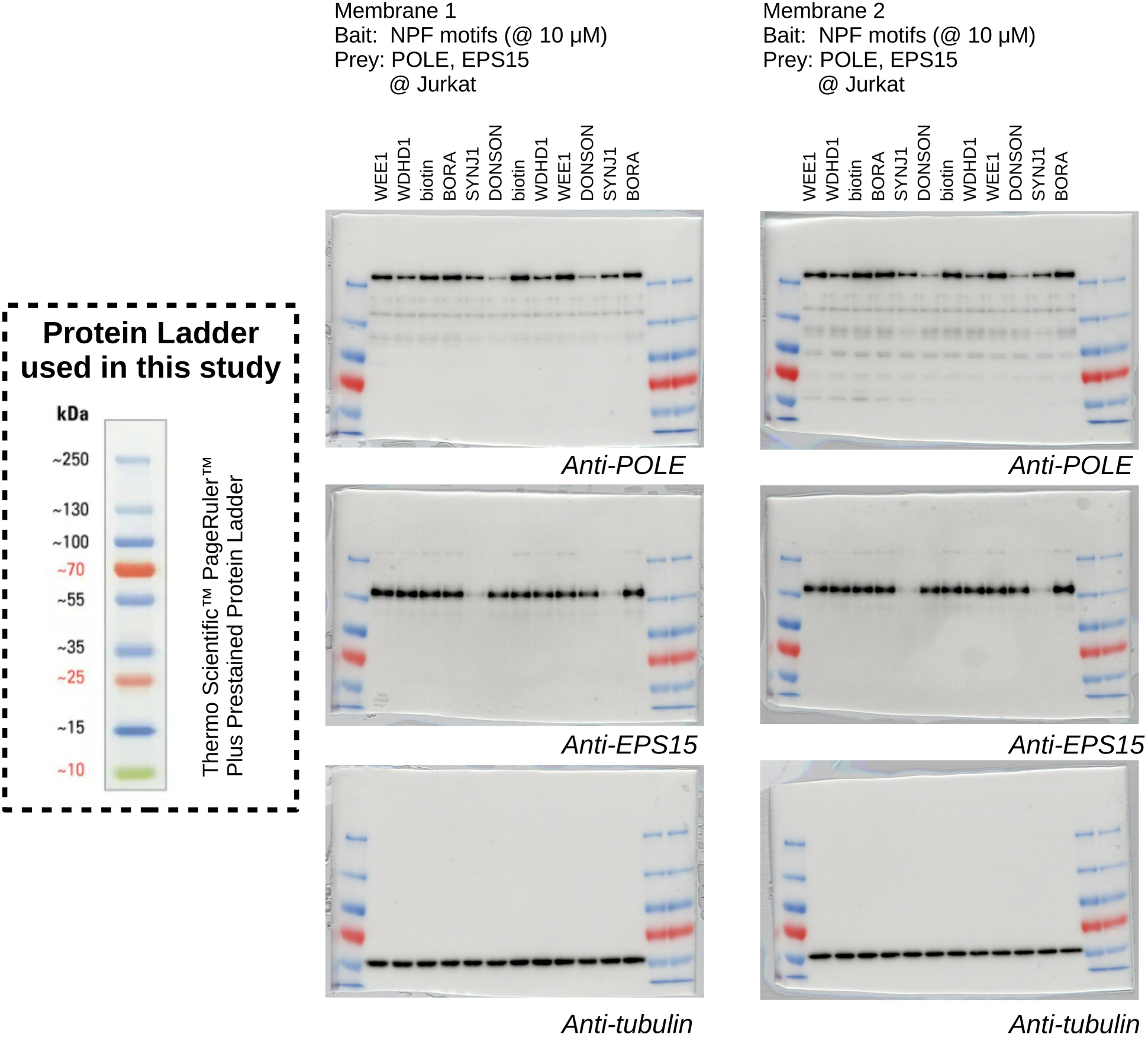
Raw data of the nHU-WB experiments shown on Figure S2. Two independent experiments are loaded on each blots and Western blots were performed in technical duplicates. Western blot images are shown as overlay of luminescence and colorimetric photos. The same protein marker is used in the entire study for which the manufacturer’s reference image is included on the left side.

**Figure S4.**
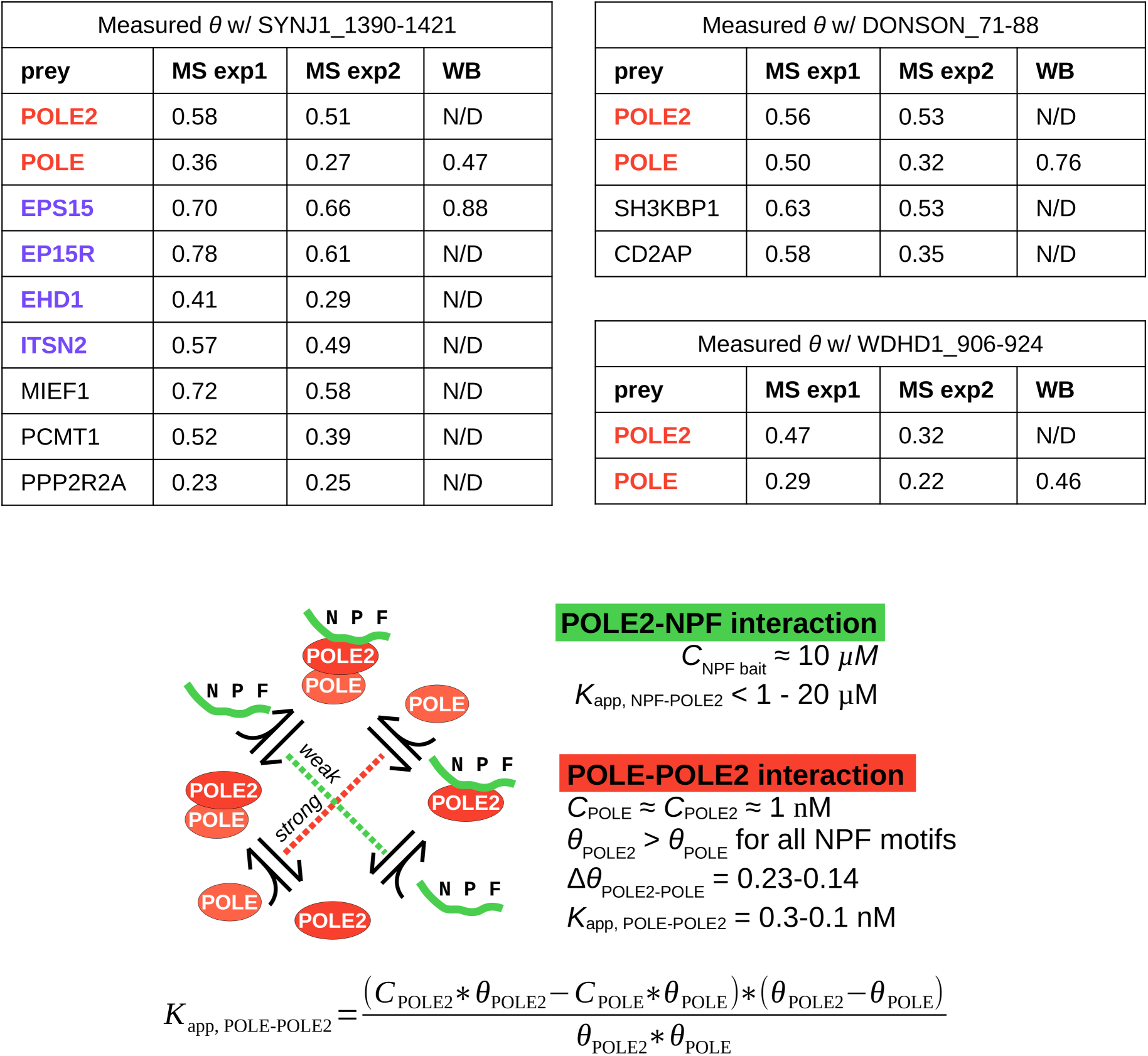
Based on apparent affinities, POLE2 is suspected to bind to NPF motifs directly. The measured degree of binding is shown for all partners that both independent experiment identified to bind to their cognate partner. In all MS experiments, the fractional depletion of POLE2 is higher than of POLE. This is consistent with a model according to which POLE2 directly binds to various NPF motifs with affinities in the ∼ μM regime, while it also binds to POLE with a dissociation constant in the ∼ nM regime.

**Figure S5.**
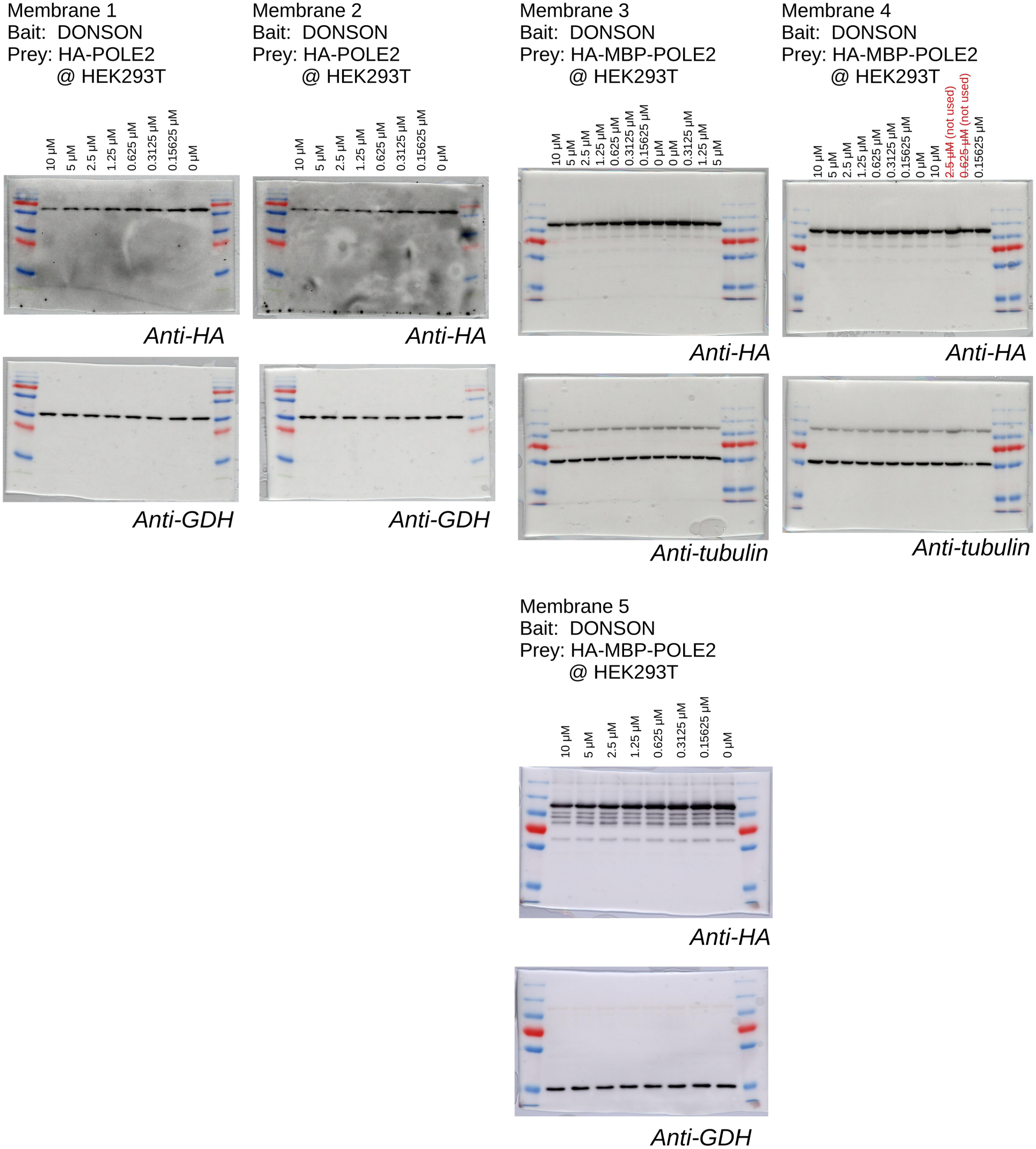
Raw data of the nHU-WB experiments shown on. Figure 1. Western blots were performed in technical duplicates or triplicates. Western blot images are shown as overlay of luminescence and colorimetric photos.

**Figure S6.**
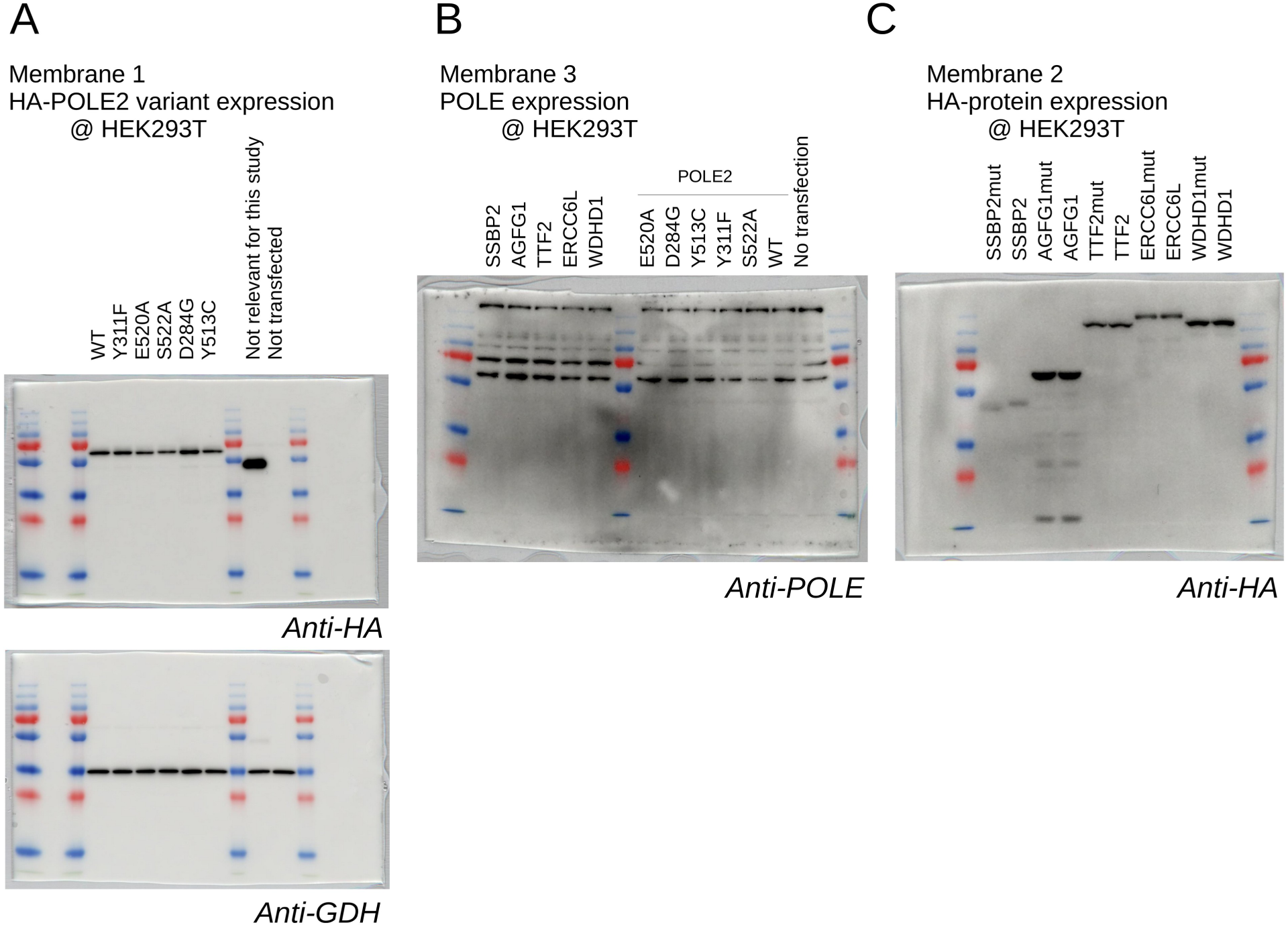
Expression analysis of transfected proteins transiently expressed in HEK293T cells. (A) Neither the expression level of POLE2 (A), POLE (B), nor its partners (C) changed substantially upon the introduction of mutations in POLE2, or NPF motifs. Western blot images are shown as overlay of luminescence and colorimetric photos.

**Figure S7.**
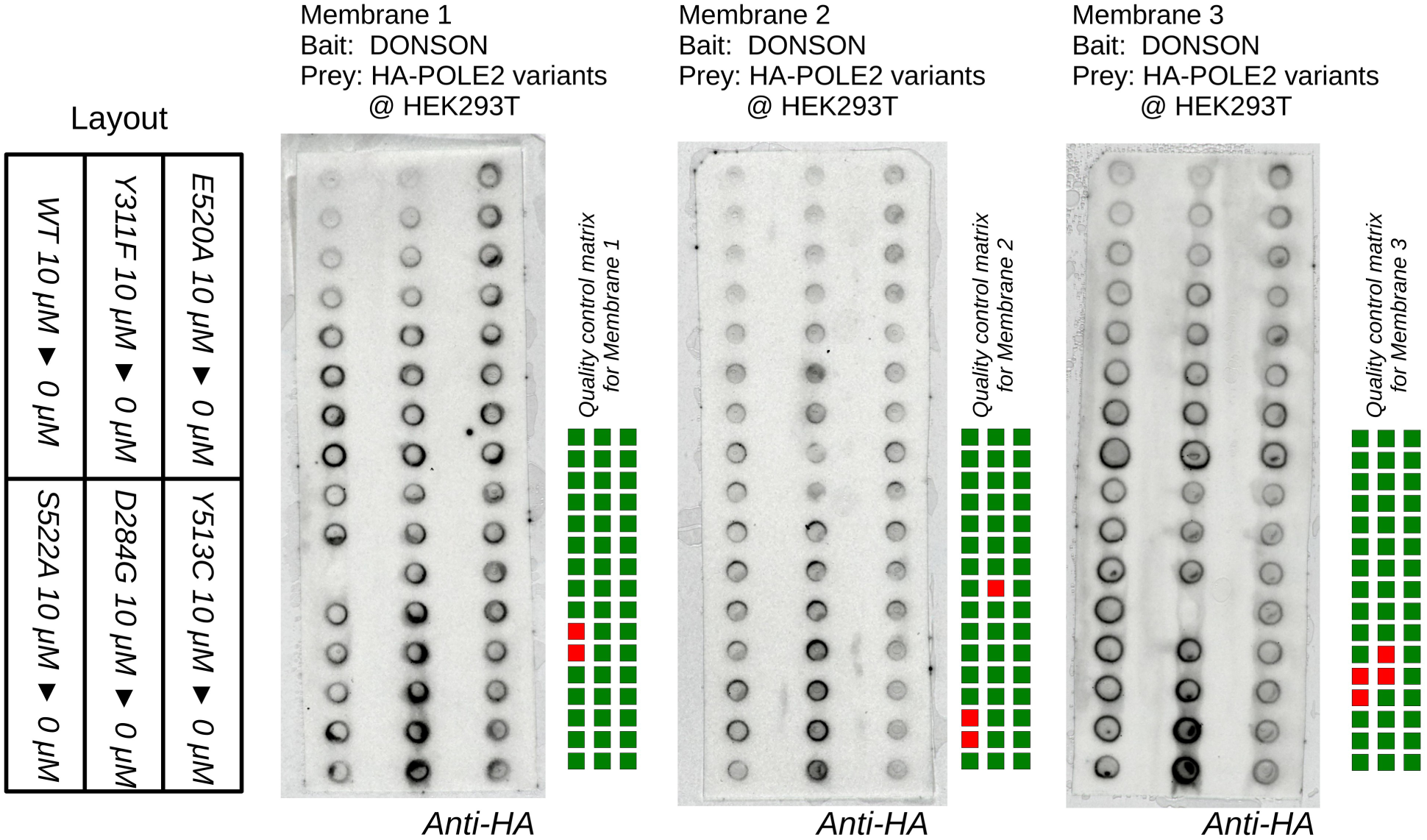
Binding of DONSON to POLE2 mutants, measured with nHU experiments coupled with dot blot analysis. On a 48 dot screen, 8-point nHU titration experiments of six POLE2 variant is included. Dot blots were performed in technical triplicates. Due to technical difficulties, such as because of complete or partial signal loss due to puncture of membrane under vacuum, specific wells were omitted from analysis. This is indicated by a quality control matrix on the right side of each membrane.

**Figure S8.**
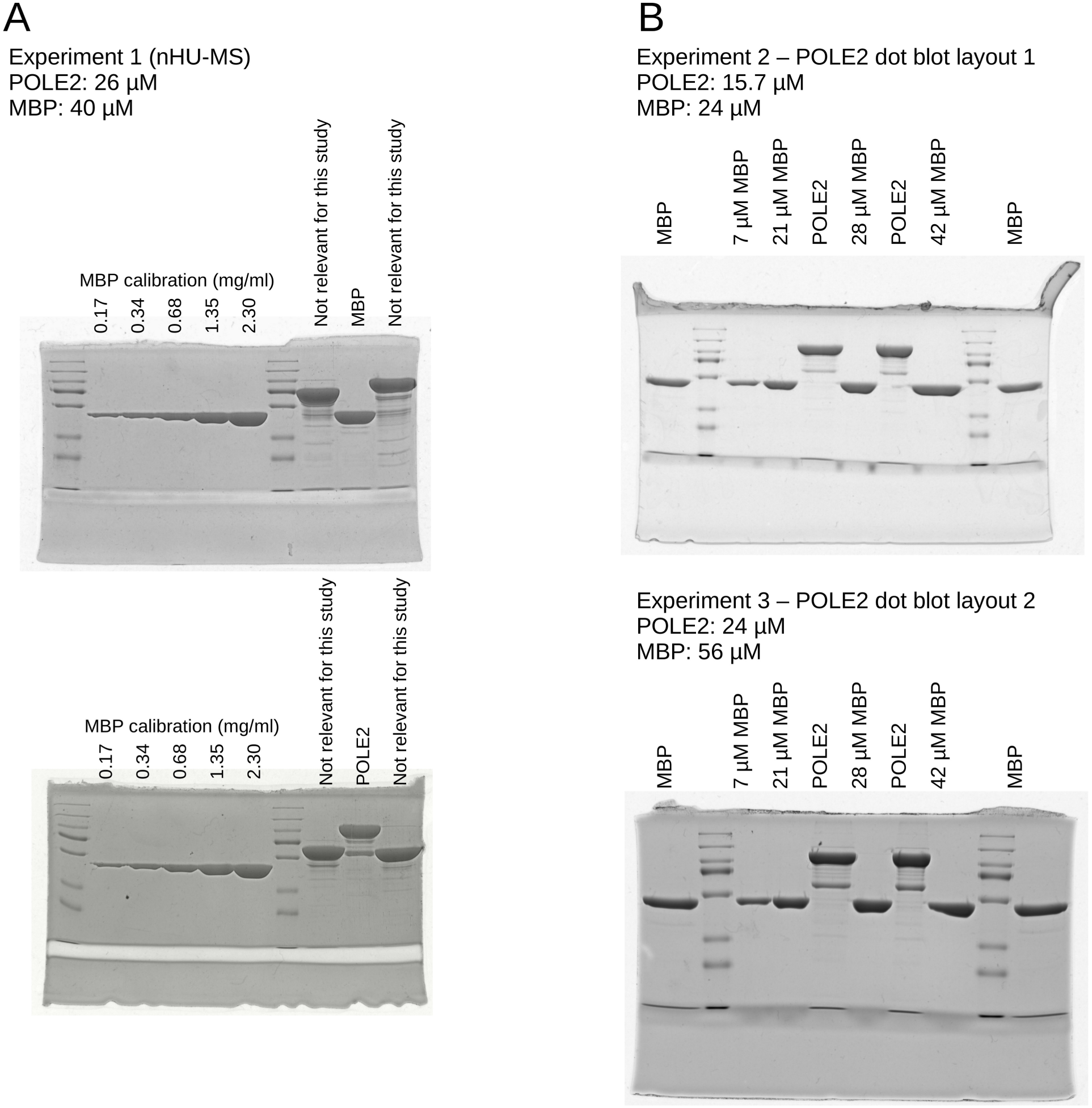
Bait concentration measurement of nHU experiments carried out with protein baits. Bait concentrations of independent nHU experiments analyzed by MS (A), or dot blot (B) were analyzed independently. On each Coomassie-stained gel, a series of known amounts of purified MBP protein is loaded as a calibration standard alongside the eluted protein samples. Based on densitometry and the known molecular weight difference between MBP and the proteins of interest, the bait concentration can be deduced.

**Figure S9.**
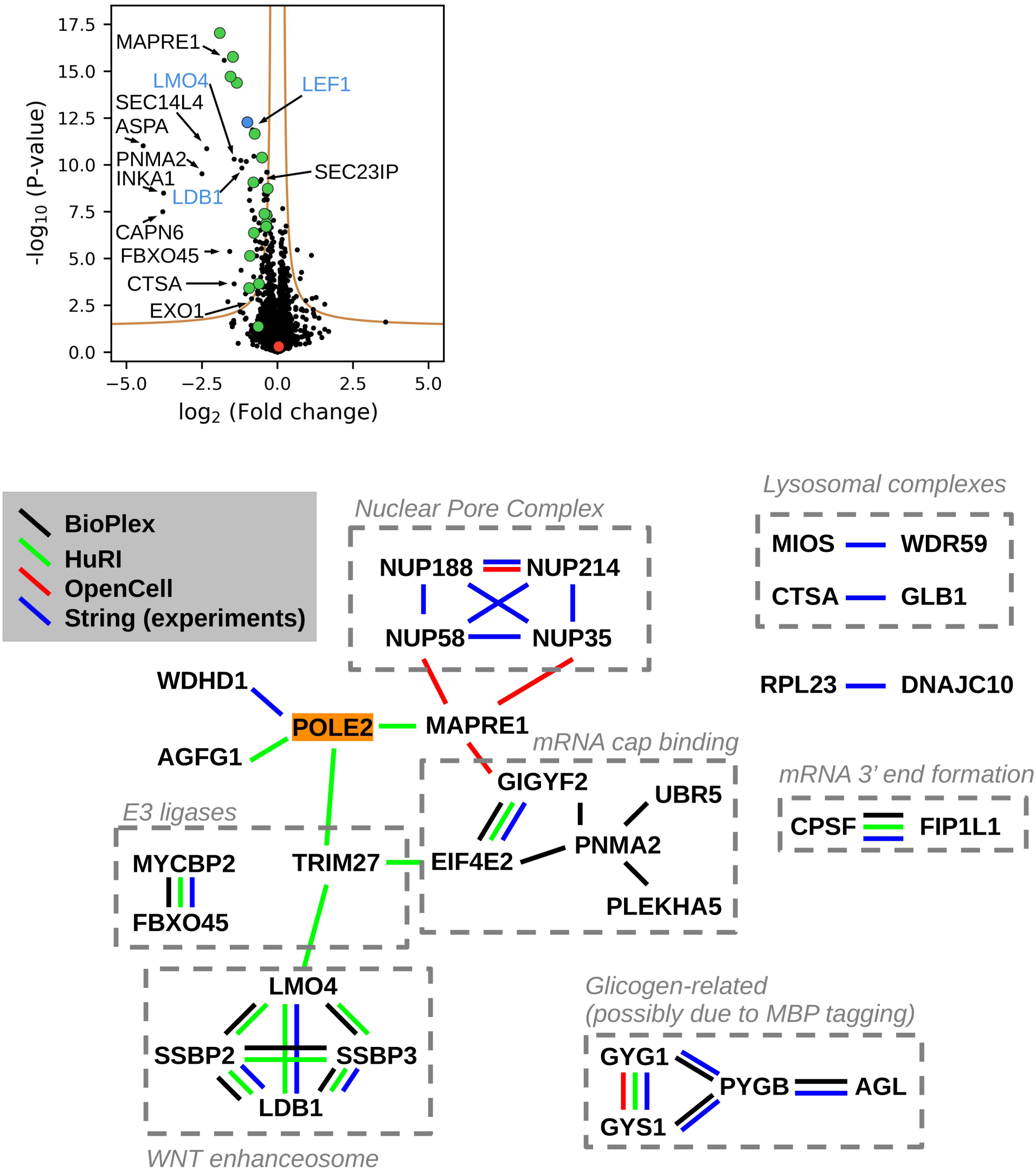
Additional details for the proteome-wide interaction network of POLE2. (top) Other notable partners identified to bind to POLE2 that could not be showed on Figure 3 for clarity. Partners marked in blue are part of the WNT-enhanceosome. (bottom) Identified complexes among POLE2 partners.

**Figure S10.**
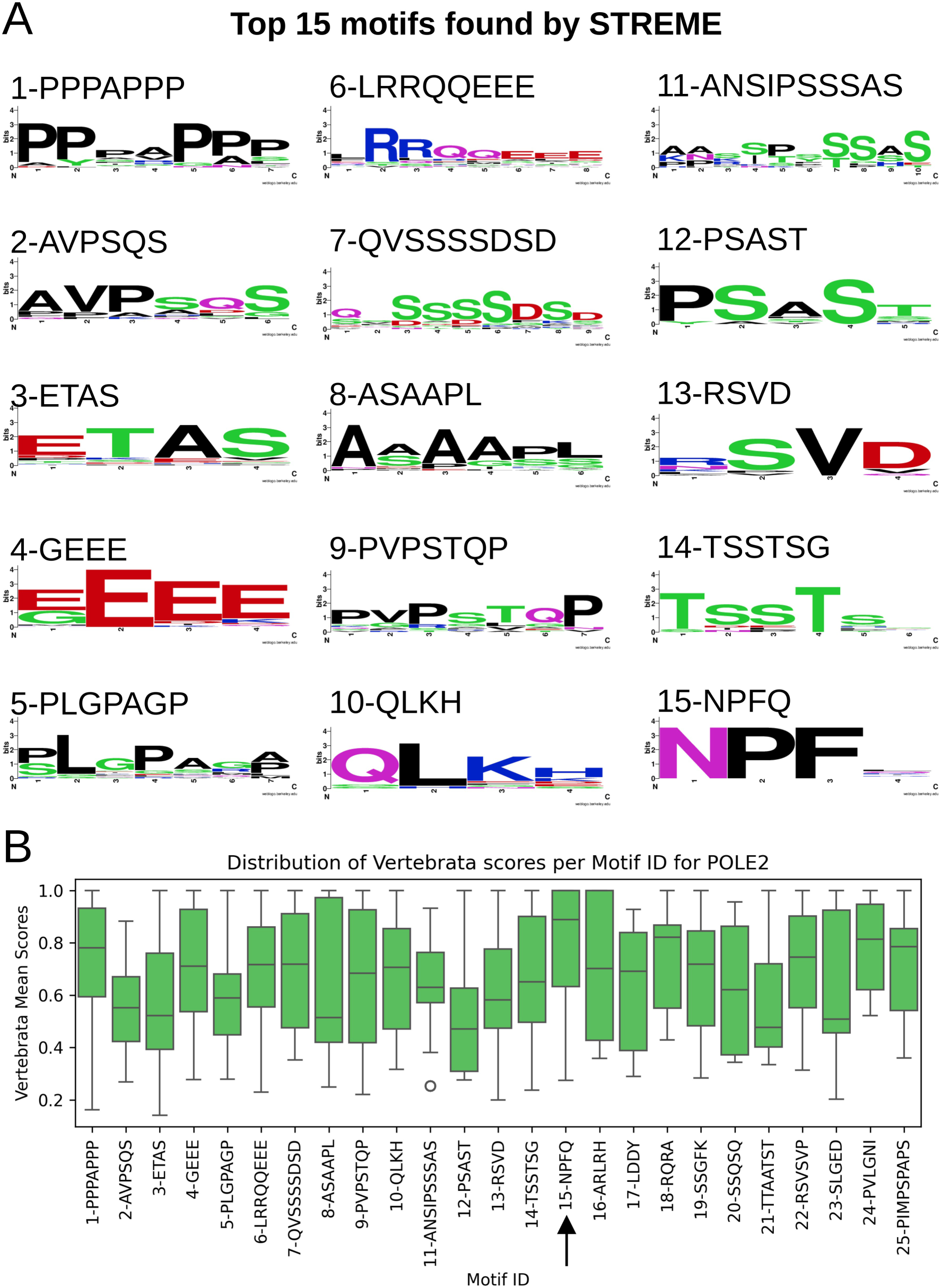
Enriched motifs in the sequences of POLE2 partners. (A) The consensus logo of the top 15 motif classes identified by STREME using the disordered sequences of the partner proteins. (B) Taking into consideration the average motif evolutionary conservation within each identified motif class, the most conserved motif class turned out to be the NPFQ motif, originally ranked as #15 most enriched motif type.

**Figure S11.**
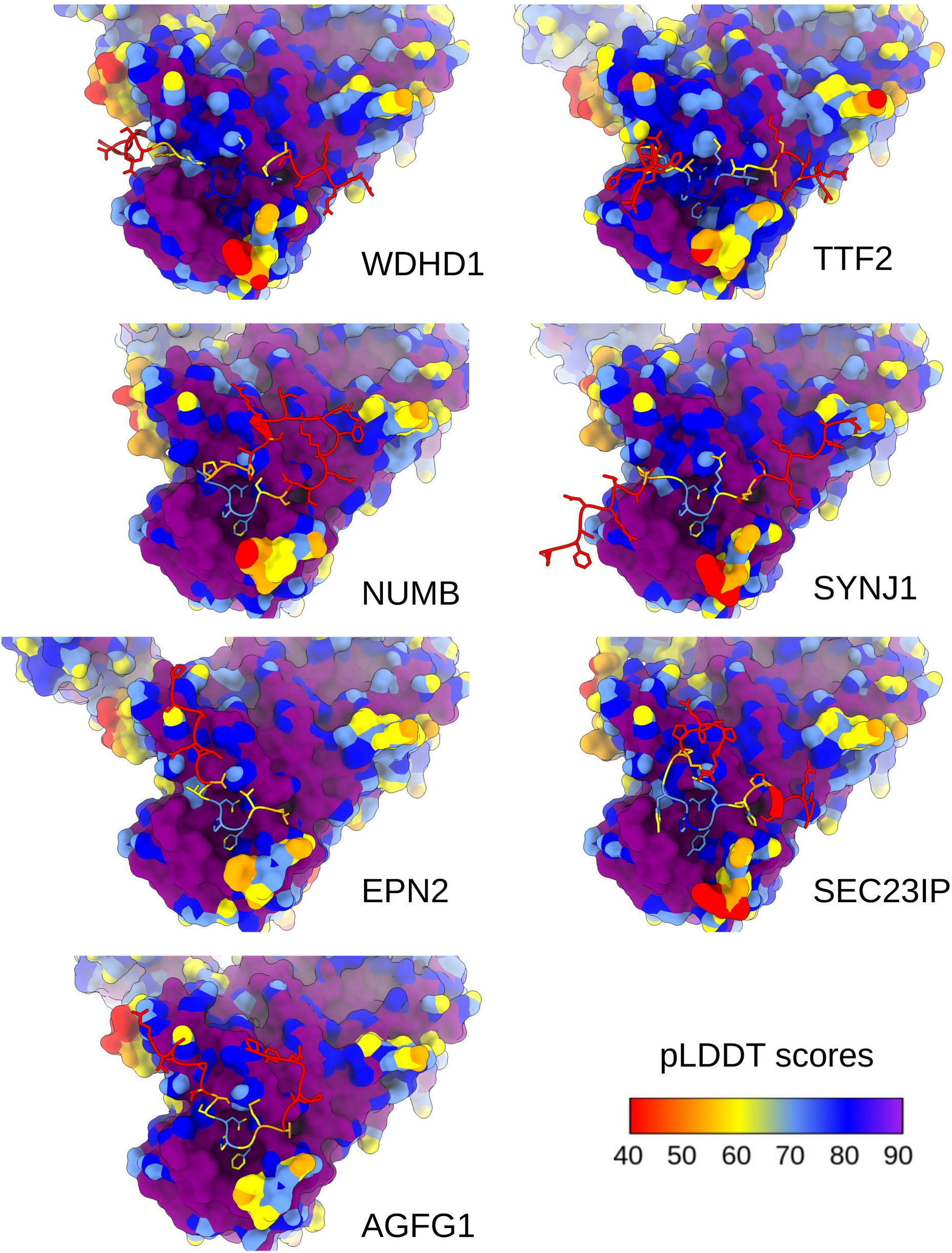
NPF motifs are predicted to bind to POLE2 with high confidence. Only the predicted NPF/NPY motifs are shown in these illustrations out of the binary AlphaFold predictions between pairs of full-length proteins. The surface of POLE2, as well as the predicted motifs of partner proteins are colored according to their pLDDT scores. High confidence is indicated by high local pLDDT scores that are outstanding in the proximity of NPF motifs and rapidly decrease in flanking sequences.

**Figure S12.**
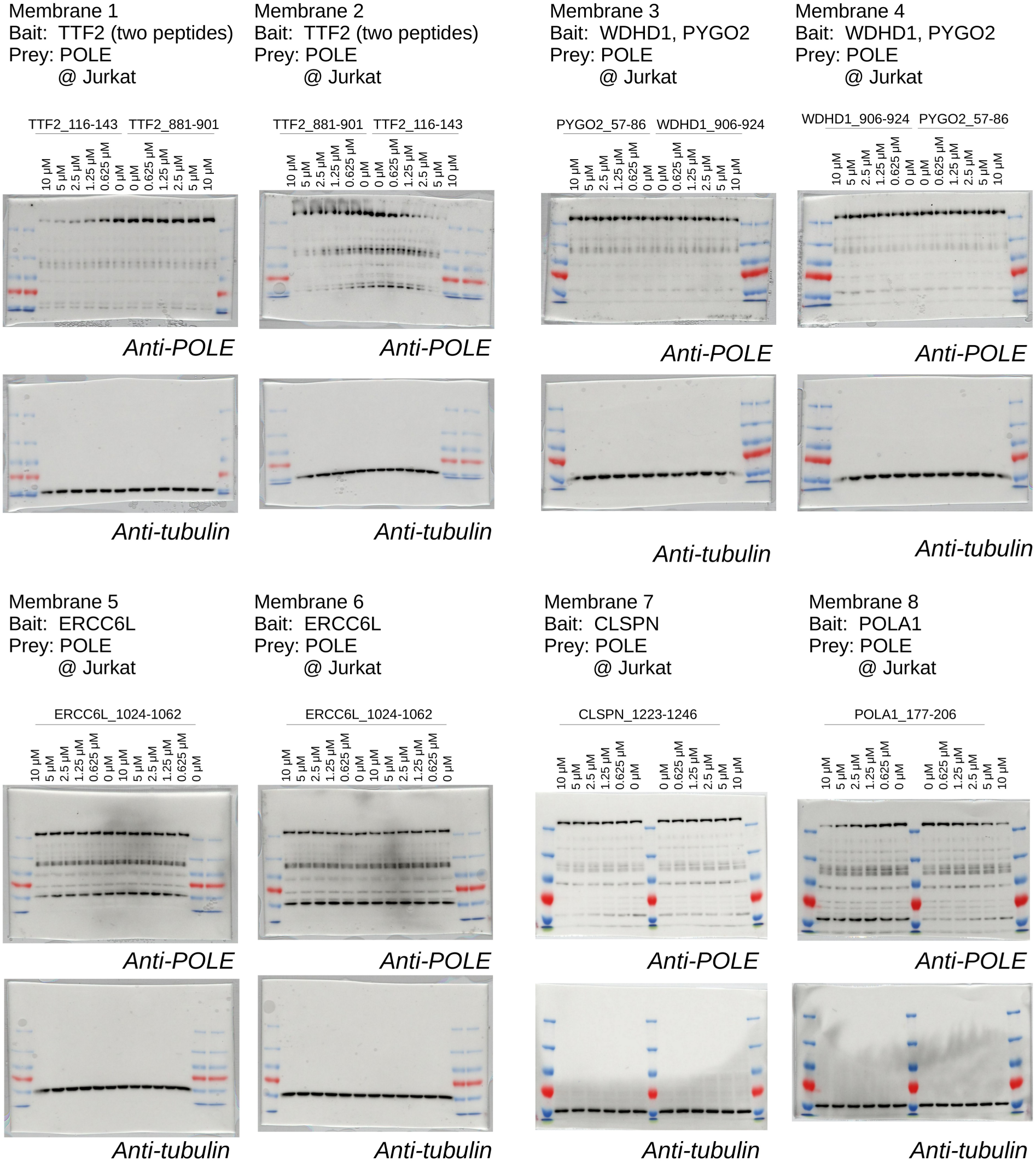
NPF motifs taken out of selected identified full-length partners bind to POLE. Titration nHU experiments with various NPF peptide baits were analyzed by Western blot to measure POLE depletion. Western blots were performed in technical duplicates. In the case of ERCC6L, the experiment was carried out in duplicates. Western blot images are shown as overlay of luminescence and colorimetric photos.

**Figure S13.**
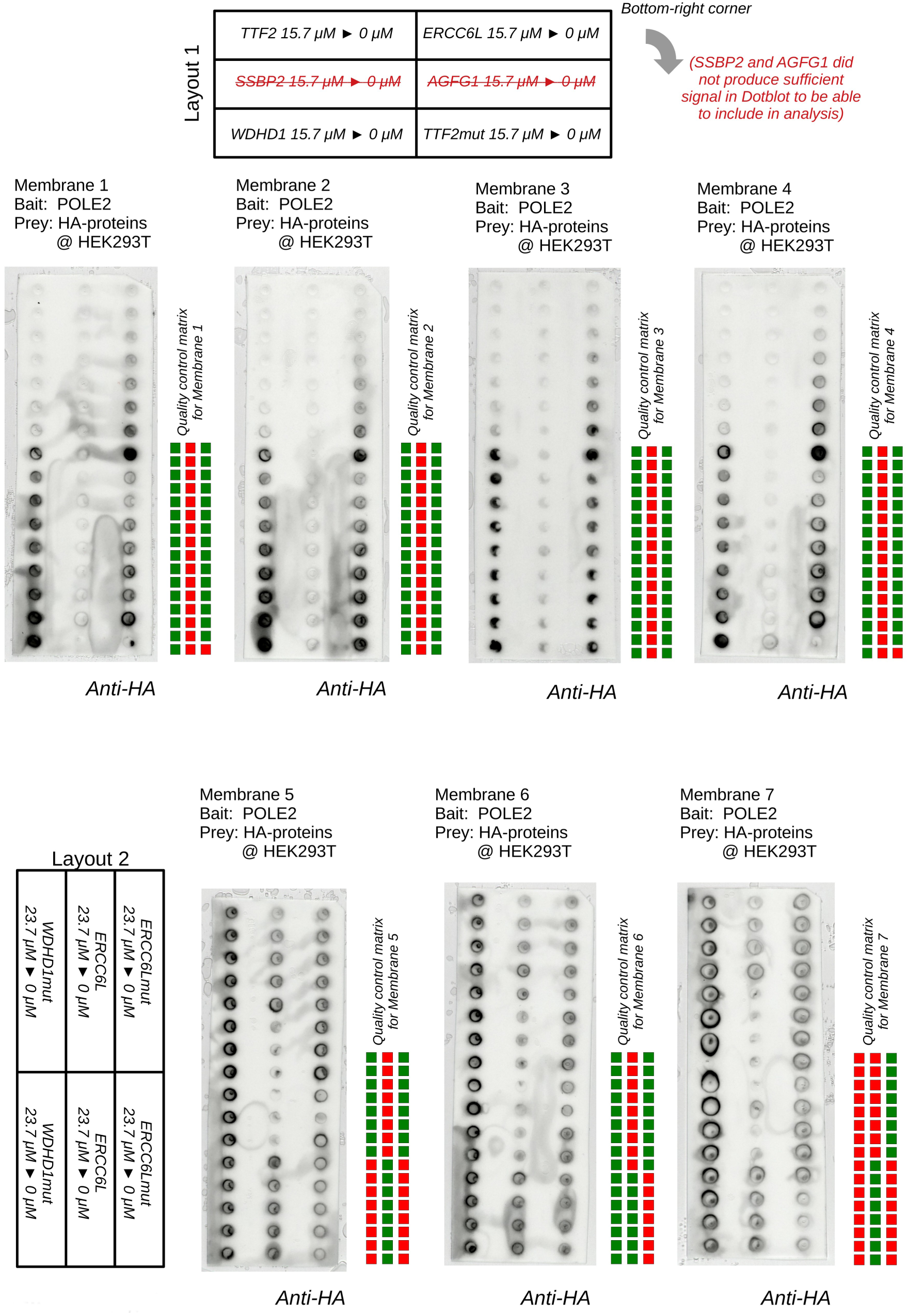
Binding of POLE2 to full-length proteins with wild type (NPF) or mutant (KAF) motifs, measured with nHU experiments coupled with dot blot analysis. On a 48 dot screen, 8-point nHU titration experiments of six protein variants are included, however some proteins were not suitable for dot blot quantification. Dot blots were performed in technical triplicates or quadriplicated. Due to technical difficulties, such as because of complete or partial signal loss due to puncture of membrane under vacuum, specific wells were omitted from analysis. This is indicated by a quality control matrix on the right side of each membrane.

**Figure S14.**
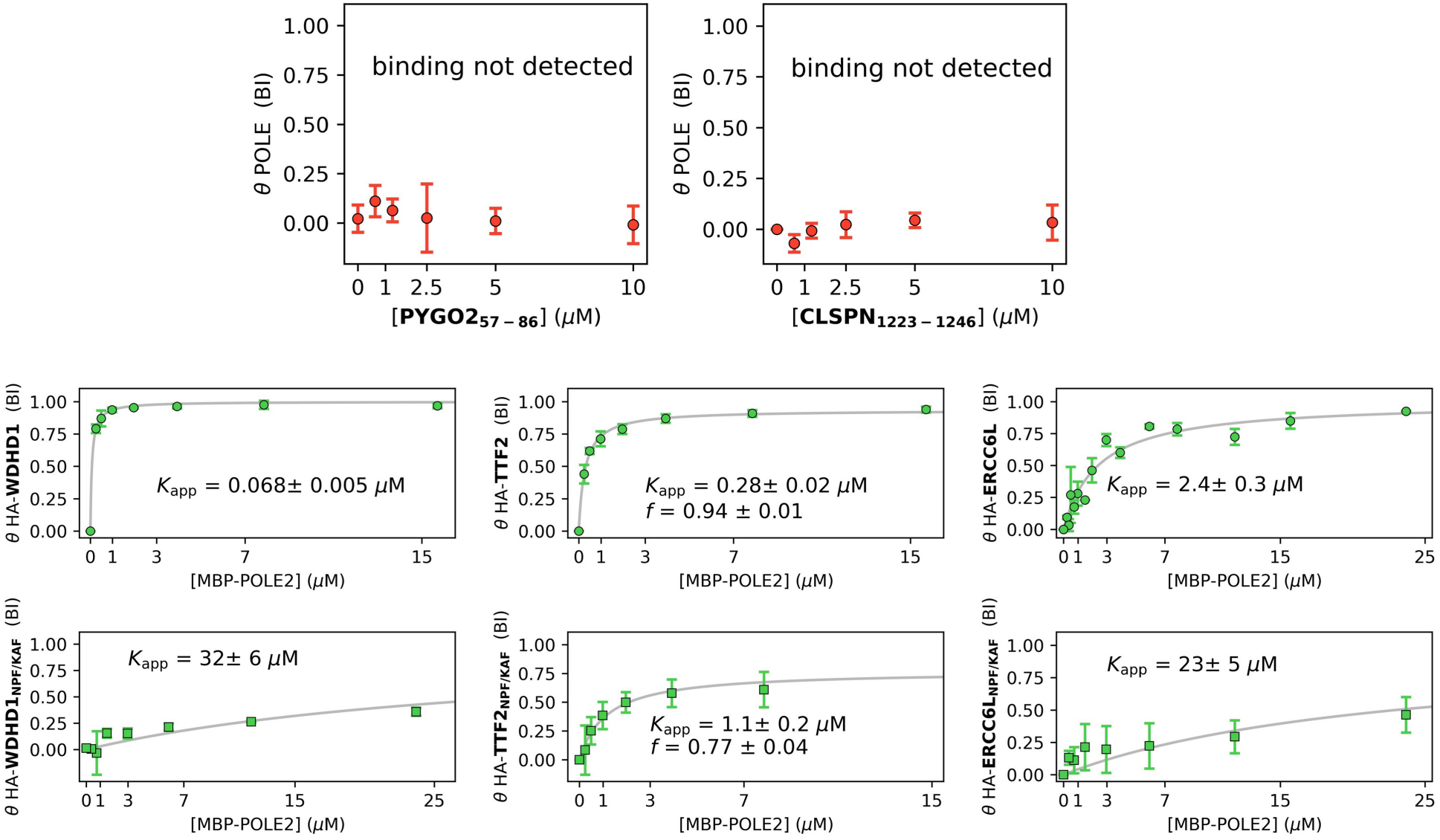
Results of nHU titration experiments. (top) The NPF motifs of PYGO2 and CLSPN were found to be unable to bind to endoganous POLE. This is consistent with the finding of the MS results shown on Figure 1, as well as on Figure 3. (bottom) Binding isothermes of POLE2 partners. Partial binding activity had to be used in the case of TTF2 interactions.

**Figure S15.**
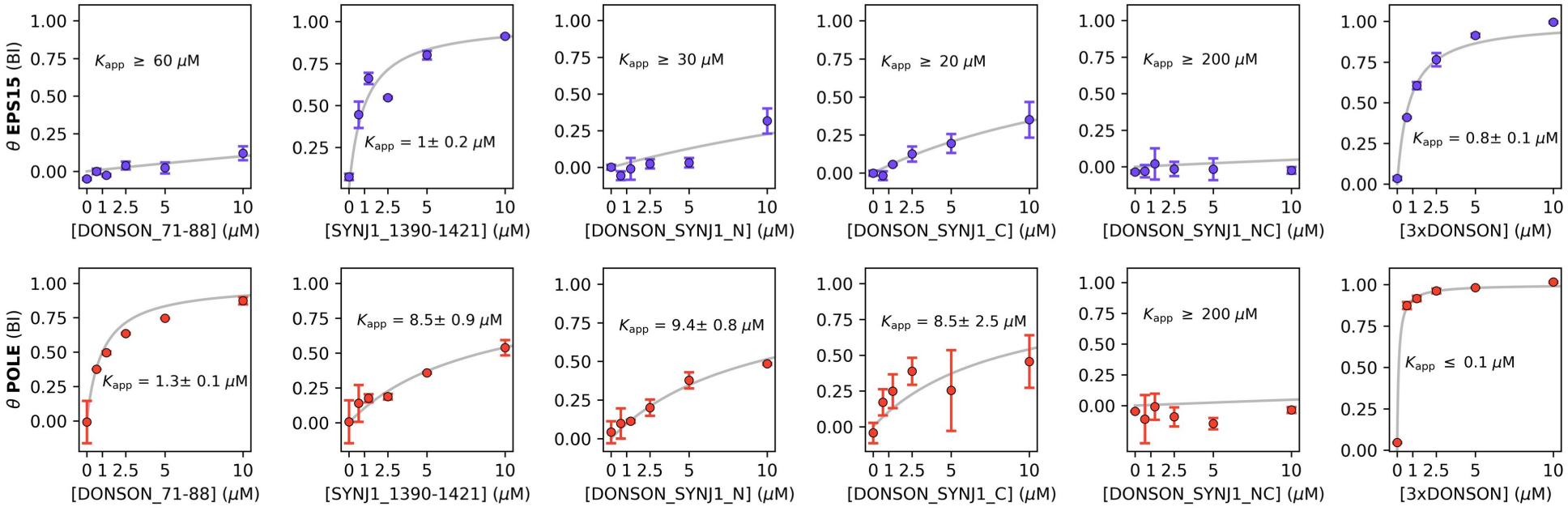
Results of nHU titration experiments with SYNJ1 and DONSON chimera NPF motif peptides. The binding of each peptide was measured against EPS15, as well as POLE.

**Figure S16.**
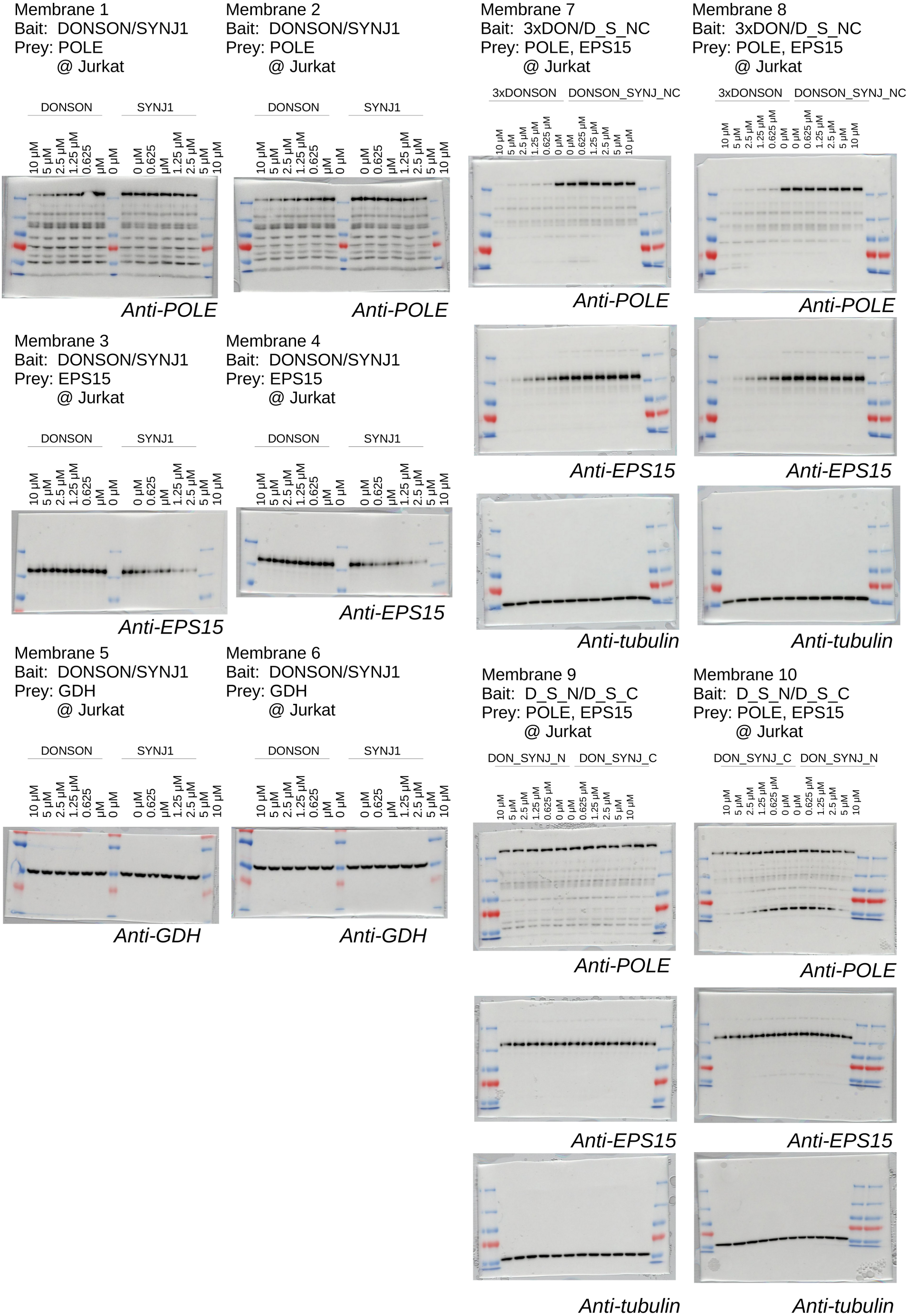
Raw data of the nHU-WB experiments shown on Figure S15. Western blots were performed in technical duplicates. Western blot images are shown as overlay of luminescence and colorimetric photos.

**Figure S17.**
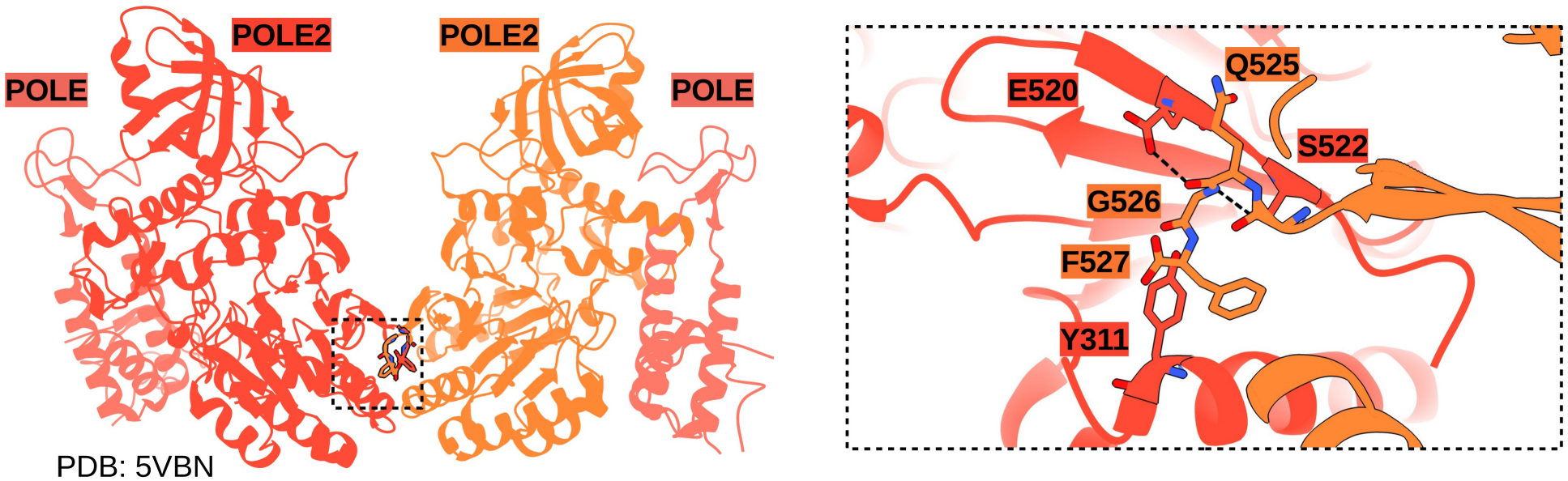
POLE2 in crystallo interacts with its own C-terminal tail through the same binding interface used to capture NPF motifs. In a previously solved crystal structure, a POLE2 dimer was observed where the non-conserved C-terminal tail of the protein (sequence: LQGF-COOH) was found to interact symmetrically with the same binding pocket used by NPF motifs, formed by S522, E520, Y311. The last Phe residue critical for such interaction is only present in mammalians and there is no evidence about in solution POLE2 dimers. Thus such interaction seems to be a crystallographic artifact, yet it is likely caused good performance of identifying NPF-mediated interactions in AlphaFold predictions and its also signifies the capacity of the binding pocket to interact with an aromatic sidechain, such as the one found in NPF or NPY motifs.

